# Small molecules reveal differential shifts in stability and protein binding for G-quadruplex RNA

**DOI:** 10.1101/2025.02.10.637408

**Authors:** Justin G. Martyr, Martina Zafferani, Morgan A. Bailey, Marek D. Zorawski, Nadeska I. Montalvan, Dhanasheel Muralidharan, Michael C. Fitzgerald, Amanda E. Hargrove

## Abstract

The potential of therapeutically targeting RNA with small molecules continues to grow yet progress is hindered by difficulties in determining specific mechanisms of action, including impacts on RNA-protein binding. RNA G-quadruplexes (rGQs) are a particularly promising target due to their range of biological functions, structural stability, and hydrophobic surfaces, which promote small molecule and protein interactions alike. Challenges arise due to 1) the low structural diversity among rGQs, thereby limiting binding selectivity, and 2) a lack of knowledge regarding how small molecules can manipulate rGQ-protein binding on a global scale. We first leveraged a small molecule library privileged for RNA tertiary structures that displayed differential binding to rGQs based on loop length, consistent with computational predictions for DNA GQs. We next utilized an RT-qPCR-based assay to measure stability against enzymatic readthrough, expected to be a common mechanism in rGQ function. We discovered small molecules with significant, bidirectional impacts on rGQ stability, even within the same scaffold. Using Stability of Proteins from Rates of Oxidation (SPROX), a stability-based proteomics method, we then elucidated proteome level impacts of both stabilizing and destabilizing rGQ-targeting molecules on rGQ-protein interactions. This technique revealed small molecule-induced impacts on a unique subset of rGQ-binding proteins, along with proteins that exhibited differential changes based on the identity of the small molecule. The domain and peptide-level insights resulting from SPROX allow for the generation of specific hypotheses for both rGQ function and small molecule modulation thereof. Taken altogether, this methodology helps bridge the gap between small molecule-RNA targeting and RNA-protein interactions, providing insight into how small molecules can influence protein binding partners through modulation of target RNA structures.

## INTRODUCTION

Small molecule targeting of RNA shows immense promise for future therapeutics.^1-3^ From clinical successes targeting the splicing machinery of SMN2 pre-mRNA^4^ and others in development, our understanding of the small molecule and RNA properties that drive *binding* has significantly improved over the last decade. However, there is a significant gap in knowledge as to how these interactions ultimately lead to changes in observed biological function. Because much of RNA function arises from its interactions with RNA-binding proteins, it is reasonable to expect that these small molecules may be modulating protein binding partners, but this has not been explored on the proteomic scale.

G-quadruplexes (GQs) are biologically important nucleic acid structures that offer a unique opportunity to reveal how small molecule:RNA binding can alter interactions between target RNA structures and protein binding partners. GQs are formed from guanine-rich DNA or RNA sequences (commonly 5’-G_≥2_N_1-7_G_≥2_N_1-7_G_≥2_N_1-7_G_≥2_ -3’)^5^ that can form planar G-quartets through Hoogsteen base-pairing interactions. These quartets can then stack in 3D space by coordinating metal cations, forming complex tertiary structures.^6-9^ Structural diversity among GQs arises primarily from differences in quartet number, loop length and composition, and topology/strand directionality, as well as slight modifications to the core structure such as single nucleotide bulges between quartets.^10-12^ However, variations among these parameters are limited, rendering GQs a specificity challenge for nucleic acid molecular recognition despite the unique 3D structure.

While initial chemical probing studies suggested cellular mechanisms actively work to unfold RNA G-quadruplexes (rGQs),^13^ mounting evidence that rGQs not only fold in cells but are integral to biological functions has renewed interest in rGQs as potential therapeutic targets.^14-17^ GQs (both RNA and DNA) have been the aim of many targeting approaches, including synthetic peptides,^18^ oligonucleotide aptamers,^19,20^ and small molecules.^21,22^ Small molecules have gained significant traction due to their use as chemical probes to identify rGQs and G-quadruplex binding proteins (GQBPs).^23-27^ Since 1997 approximately 3000 GQ-binding small molecules have been identified, largely for DNA GQs.^28,29^ Some classic examples of these small molecules include pyridostatin (PDS) and TMPyP4 which have been used across multiple studies to validate GQ folding *in vitro*.^30, 31^ Groups have started to gain success in identifying small molecules with selectivity between rGQs and their DNA counterparts through implementation of high-throughput small molecule screening. For example, two groups separately targeted and stabilized an rGQ in the 5’-UTR of the NRAS mRNA in order to prevent ribosome readthrough without impacting *NRAS* transcription.^32,33^

While initially characterized by the inhibition of enzymatic readthrough, the known functions of rGQs have expanded drastically over recent years. They have been associated with regulating mRNA translation,^34^ modulating alternative splicing,^35^ facilitating membraneless phase separation,^36,37^ and impacting protein stability.^38^ rGQs have also been implicated in disease, including dysregulation in cancer^39^ and in neurodegenerative disease.^40-42^ Several recent efforts have sought to identify GQBPs that may be responsible for these functions.^43-45^ Approaches to identify GQBPs have focused mainly on affinity-based pulldown methods^27,46-48^ and specific interactions are often validated by varying buffer cations to impact GQ folding or by employing sequence mutations.^36,46,49,50^

Due to these efforts, there has been a growing appreciation for the impact that rGQ folding dynamics have on rGQ-GQBP interactions and downstream biological function. For example, helicases (e.g., DEAH-box helicase 36 (DHX36)) have been linked to unfolding rGQs in the 5’-UTRs of mRNAs, allowing for dynamic regulation of translation.^51,52^ Folded and unfolded rGQs may also serve as ‘landing pads’ for hnRNPF and hnRNPH1 in order to scaffold splicing machinery and promote both exon inclusion and exclusion.^53-55^ As of yet, limited studies have examined mechanisms by which small molecules may similarly encourage the folding and unfolding of rGQs, with a few focused on evaluating small molecule properties that drive differential stabilization of these structures,^30,56^ and others using pan-rGQ binding small molecules to manipulate rGQ-GQBP interactions.^49,54, 57^ In one compelling example, Caplen and co-workers reported that targeting an rGQ at the hnRNPH1-dependent *EWSR1* splice site with a pan-GQ stabilizer decreased hnRNPH1 binding and effectively inhibited fusion splicing of *EWS-FLI1*, with the authors emphasizing the importance of evaluating the specific interactions involved to understand this biological readout in future work.^54^ Revealing the impacts of RNA-binding small molecules on rGQ-GQBP interactions on the proteomic scale would be a critical step forward in our mechanistic understanding of small molecule-induced changes in RNA-protein interactions and subsequent biological function.

Our group has a long standing interest in targeting the long noncoding RNA metastasis-associated lung adenocarcinoma transcript 1 (MALAT1), a transcript associated with cancer among other diseases.^58-61^ Recently, two groups identified and validated conserved rGQs in the 3’-end of MALAT1 (**Figure 1**). Kwok and co-workers targeted these MALAT1 rGQs *in vitro* and in cells with a pan-GQ ligand, a synthetic peptide, or an L-RNA aptamer, respectively, and observed displacement of two identified rGQ-GQBP interactions, specifically with DHX36 and NONO.^49^ Meanwhile, Maiti and co-workers took a phenotypic approach and found that while deletion of the same rGQs did not impact MALAT1 stability, localization, or nuclear speckle formation, it did impact localization of known GQBPs nucleolin and nucleophosmin, and binding to these proteins was confirmed through pull-down mass spectrometry.^50^ These studies inspire further investigations into how these MALAT1 rGQs may function in disease or healthy biology and therefore represent a promising set of rGQs for small molecule targeting in tandem to evaluating rGQ-GQBP modulation.

**Figure 1.**
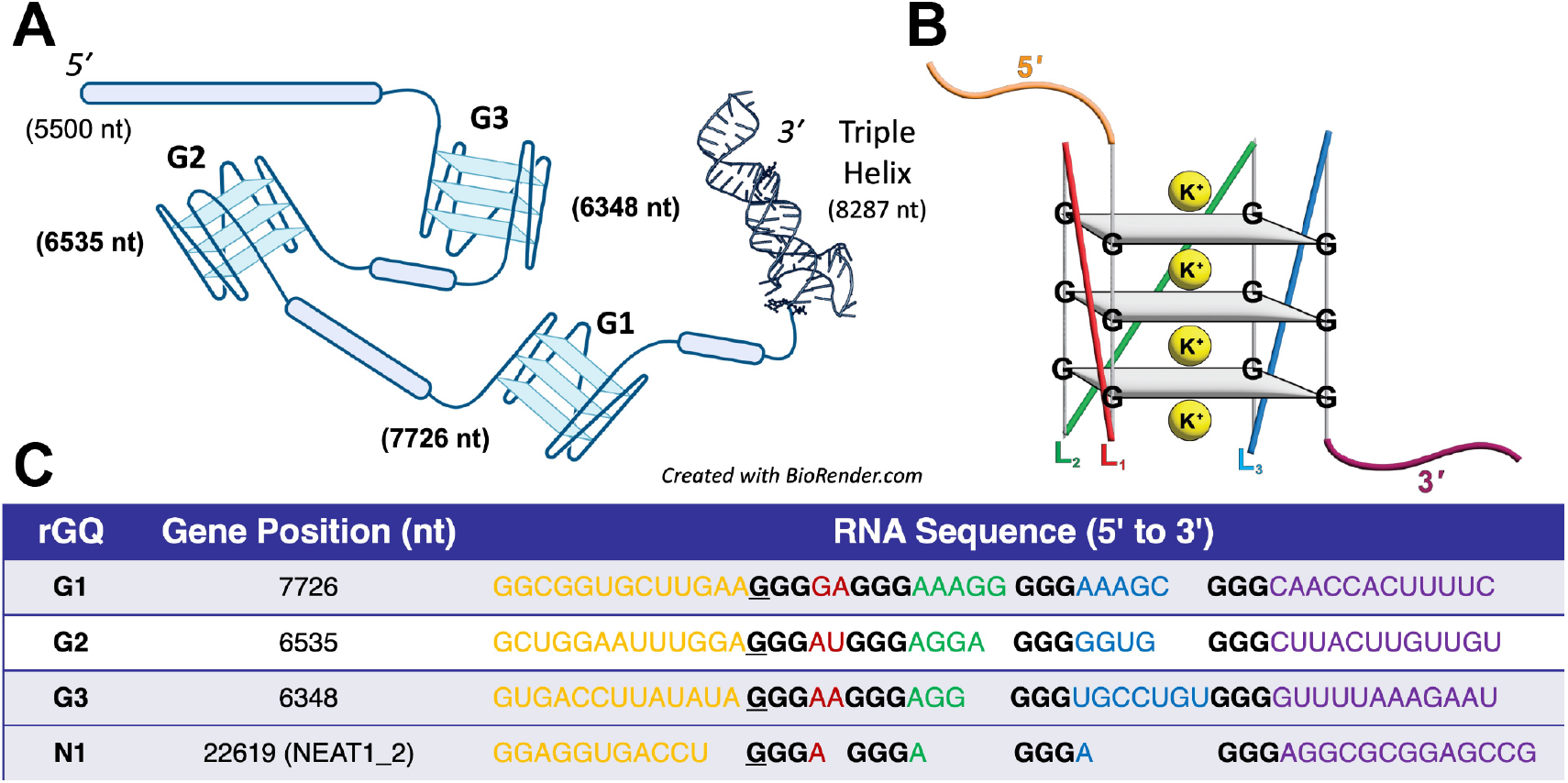
A) Schematic of MALAT1 rGQ placement and nucleotide position in the MALAT1 gene sequence (NCBI: NC_000011.10) with the 3’ triple helix (TH) also annotated. Created with BioRender.com. B) Parallel topology G-quadruplex structure with G-tetrads, loops (L_1_, L_2_, L_3_) and 5’ and 3’ flanking regions distinguished by color, C) MALAT1 and NEAT1 rGQ sequences used in this study with sequences colored to reflect the G-quadruplex structure (G-tracts in black, 5’ flanking sequence in orange, L_1_ in red, L_2_ in green, L_3_ in blue, and 3’ flanking sequence in purple).

In this study, we utilized an in-house synthetic library of diminazene (DMZ) molecules that had been previously shown to differentially modulate stability of another tertiary stucture, namely the 3’-end MALAT1 triple helix (TH).^60^ DMZ molecules have also been found to bind DNA GQs with low-micromolar affinity^62,63^ and computationally predicted to bind DNA GQs with differing affinities based on the GQ loop length.^64^ Based on this precendent, we assessed differential binding to three conserved MALAT1 rGQs and evaluated the ability of these molecules to impact rGQ stability and folding. We further employed a stability-based proteomic method, namely Stability of Proteins from Rates of Oxidation (SPROX), to understand if and how these DMZ molecules modulate rGQ interactions with proteins, providing the first case study to evaluate how small molecules can differentiate between rGQs, modulate rGQ stability, and impact rGQ-protein interactions on the proteomic scale.

## RESULTS AND DISCUSSION

### Diminazene screening and affinity for rGQs

Prior to previous reports,^49,50^ we independently identified the three well-conserved MALAT1 rGQs, hereafter referred to as **G1** (nt. 7726), **G2** (nt. 6535) and **G3** (nt. 6348) (**Figure 1**) and carried out CD and ThT validation (**Figure S1-2**), confirming their *in vitro* folding. We also selected an rGQ from previous work on NEAT1 (N1_22619, referred to as **N1**)^36^ as an rGQ control due to its well-characterized K^+^ dependence. Initial small molecule screening was performed with an in-house small molecule library designed around the diminazene (DMZ) scaffold. This scaffold has been reported to engage DNA GQs of various topologies.^62-64^ However, this scaffold has not yet been explored for rGQ binding. In previous work, our laboratory synthesized a 21-member DMZ focused small molecule library to target the MALAT1 triple helix (TH).^60^ We saw this tertiary-structure biased library as a unique opportunity to evaluate small molecule interactions across multiple RNA tertiary structures and provide a starting point for measuring small molecule-induced modulation of these different structures.

#### Indicator displacement assay screening

We utilized indicator displacement assays (IDAs) as a preliminary measure to obtain a binding affinity for each DMZ library member using the constructs described above.^65^ IDA is an established and reliable method that can be performed in a high throughput manner and requires a low amount of material. For our experiments, we used the dye RiboGreen™ as our indicator.^66^ All 21 DMZ library members were assayed at five concentrations (1, 5, 10, 25, 50 µM), including **DMZ-P0**, the parent DMZ scaffold found to bind to DNA GQs.^63^ We identified 8 molecules across our four rGQ targets with dose-dependent indicator displacement and apparent sub micromolar affinity for at least one rGQ from the initial 6-point quick screen. We then ran these 8 molecules in 10-point titrations to more finely determine their apparent affinity for each rGQ, yielding the heatmap shown in **Figure 2A** (**Figure S3**).

**Figure 2.**
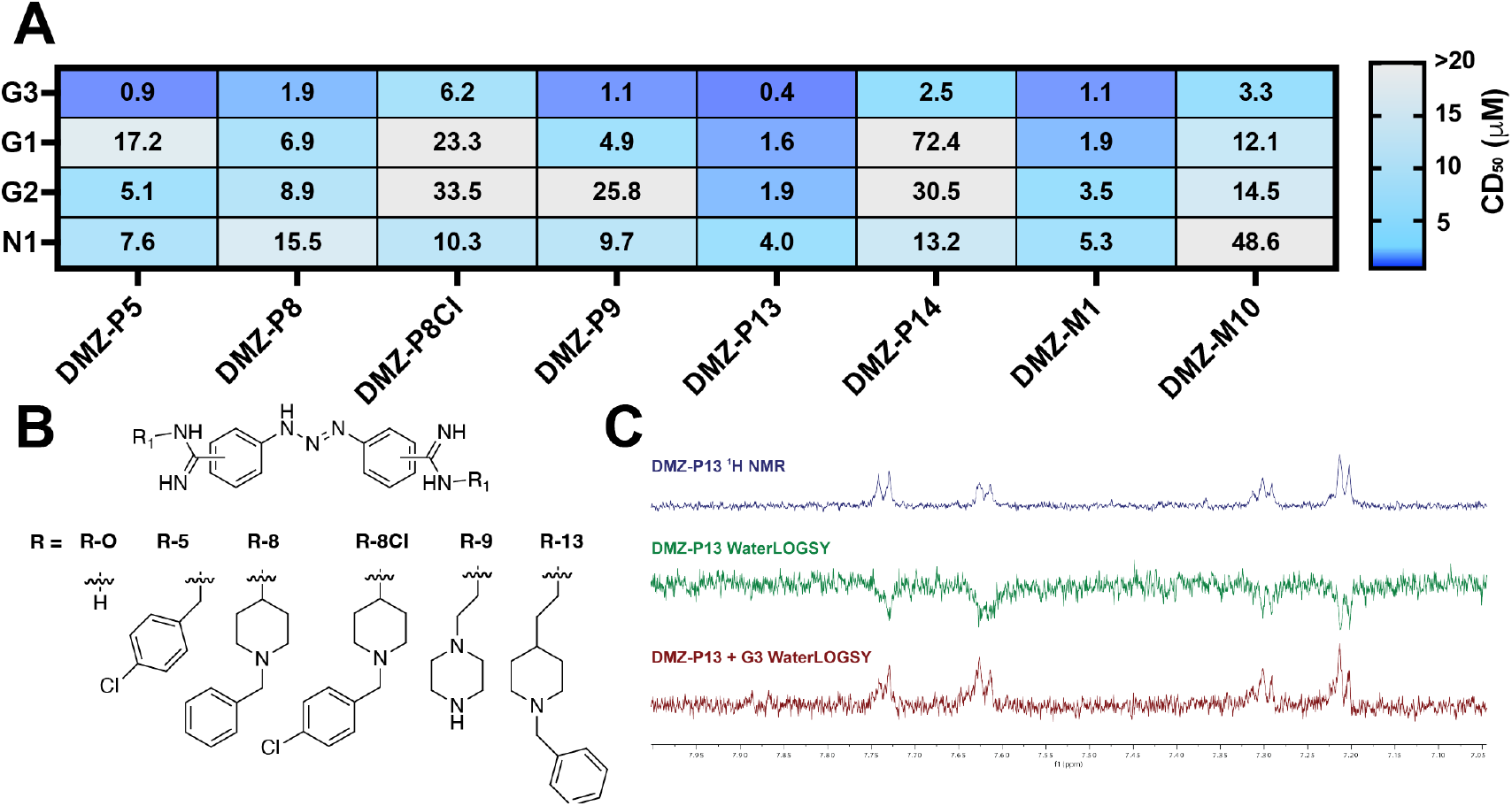
A) IDA heatmap demonstrating DMZ binding to the different rGQs, CD_50_ values shown are the mean of three independent replicates comprised of technical triplicates. B) DMZ scaffold and substituents published in previous work^60^ that demonstrate structure activity relationship trends. C) 1 mM of **DMZ-P13** was evaluated in NMR buffer (50 mM d11-Tris-HCl, 50 mM KCl, 5% DMSO pH 6.7 in 90:10 RNase-free water:D_2_O) in the presence and absence of 8 µM **G3**. Inversion of aromatic protons upon **G3** addition are shown (7.2 - 7.8 ppm).

Through this assay, we identified DMZ molecules that can bind to and displace the dye from rGQ structures with CD_50_ values in the low to sub-micromolar range. These small molecules exhibit differential binding between **G1, G2**, and **G3** as well as preferential binding for the MALAT1 rGQs over the **N1** rGQ. Comparing these CD_50_ values to those previously determined for the MALAT1 TH, the DMZs overall have higher apparent affinity for the rGQ structures than the TH.^60^ This is highlighted by **DMZ-P13**, which had approximately 7-fold higher affinity for **G3** over the TH. Conversely, TH binders **DMZ-P0** and **DMZ-P17** did not qualify as hits for the rGQs, lacking any indicator displacement, suggesting selectivity among the tertiary structures.^60^

Within the DMZ library, **DMZ-P8** and **DMZ-P13** and a new molecule synthesized for hit-to-lead TH experiments, **DMZ-P8Cl**, demonstrated structure-activity relationships that highlight the impact of these substituents on rGQ binding. The structures of these small molecules are shown in **Figure 2B. DMZ-P8** and **DMZ-P8Cl** differ by a single chloride atom para-substituted on the substituent benzene ring. The chloride addition led to a dramatic decrease in affinity for **G3**, suggesting that shifts in the distribution of electron density on the aryl groups impact rGQ recognition. Increasing the flexibility of the R-group with the addition of a 2-carbon linker (i.e. **DMZ-P8** vs **DMZ-P13**) led to a 2-fold increase in affinity for **G3**. These flexibility trends are also observed for **G1, G2**, and **N1**, suggesting that increased flexibility in the R-group may allow sampling of small molecule conformations more apt for rGQ binding. This binding data is the first evidence that DMZ-derived small molecules can bind to rGQs and identified **G3** as a primary target of interest with apparent DMZ affinities reaching the high nanomolar range.

*WaterLOGSY*. We complemented the above IDA approach with WaterLOGSY, an orthogonal, NMR-based method to directly observe RNA-small molecule interactions^67,68^ that has been recently applied to rGQ-small molecule interactions.^33^ Specifically, WaterLOGSY provides magnetization/NOE evidence by peak inversion for direct small molecule interactions in the presence and absence of a biomolecule. We tested the top hit **DMZ-P13** alone and in the presence of **G3**. We first ran a ^1^H NMR for **DMZ-P13** to test whether the small molecule aggregated in aqueous buffer (**Figure 2C**, *top*), which was ruled out by the inversion of the peaks when WaterLOGSY was applied (**Figure 2C**, *middle*). Upon the addition of the **G3** construct, we observe a full inversion of the aromatic proton peaks for **DMZ-P13** (**Figure 2C**, *bottom*), consistent with target engagement of all aromatic rings. The water peak was used to control for peak inversion and was not impacted by **G3** addition (**Figure S4**). By demonstrating direct DMZ binding to the MALAT1 rGQ constructs, WaterLOGSY validated the indirect measurements of IDA.

### Diminazene modulates rGQ stability

While IDA identified strong, nanomolar binders for the MALAT1 **G3**, it has been shown that binding affinity for RNA structures does not always correlate to modulation in structure or biological activity.^58,59^ Traditionally, the study of rGQ stability has been facilitated by circular dichroism titrations and melting temperature shift assays, along with a few examples of reverse-transcriptase stop approaches. We evaluated our DMZ-rGQ interactions across these techniques to biophysically characterize how these small molecules impact GQ stability.

#### Circular Dichroism

First, we employed CD spectroscopy similarly to the validation as above with TMPyP4, where changes in CD signal support binding and conformational rearrangement.^30^ An apparent affinity can also be determined by monitoring the CD signal at 268 nm with increasing small molecule concentration, providing a secondary validation for rGQ preferential binding.^69^ Previous titrations of the parent DMZ molecule to DNA G-quadruplexes in CD experiments demonstrated decreases in the magnitude of the 268 nm peak, suggesting destabilization.^63^

Based on the IDA experiments, we selected **DMZ-P5** and **DMZ-P13** for CD titrations with **G1, G2**, and **G3. DMZ-P13** titrations with **G1, G2**, and **G3** demonstrated dose-dependent decreases in the CD spectra as well as a slight right shift, suggesting a conformational change/shift upon small molecule binding and potentially destabilization (**Figure 3A-C**). Additionally, upon plotting and curve-fitting the normalized ellipticity at 268 nm, we were able to recapitulate the affinity trends previously observed through the IDA experiments (**Figure 3D**). Specifically, **G1** and **G2** showed more right-shifted curves and decreased changes in magnitude when compared to **G3**, which approached saturation at 50 µM (**Figure S5**).

**Figure 3.**
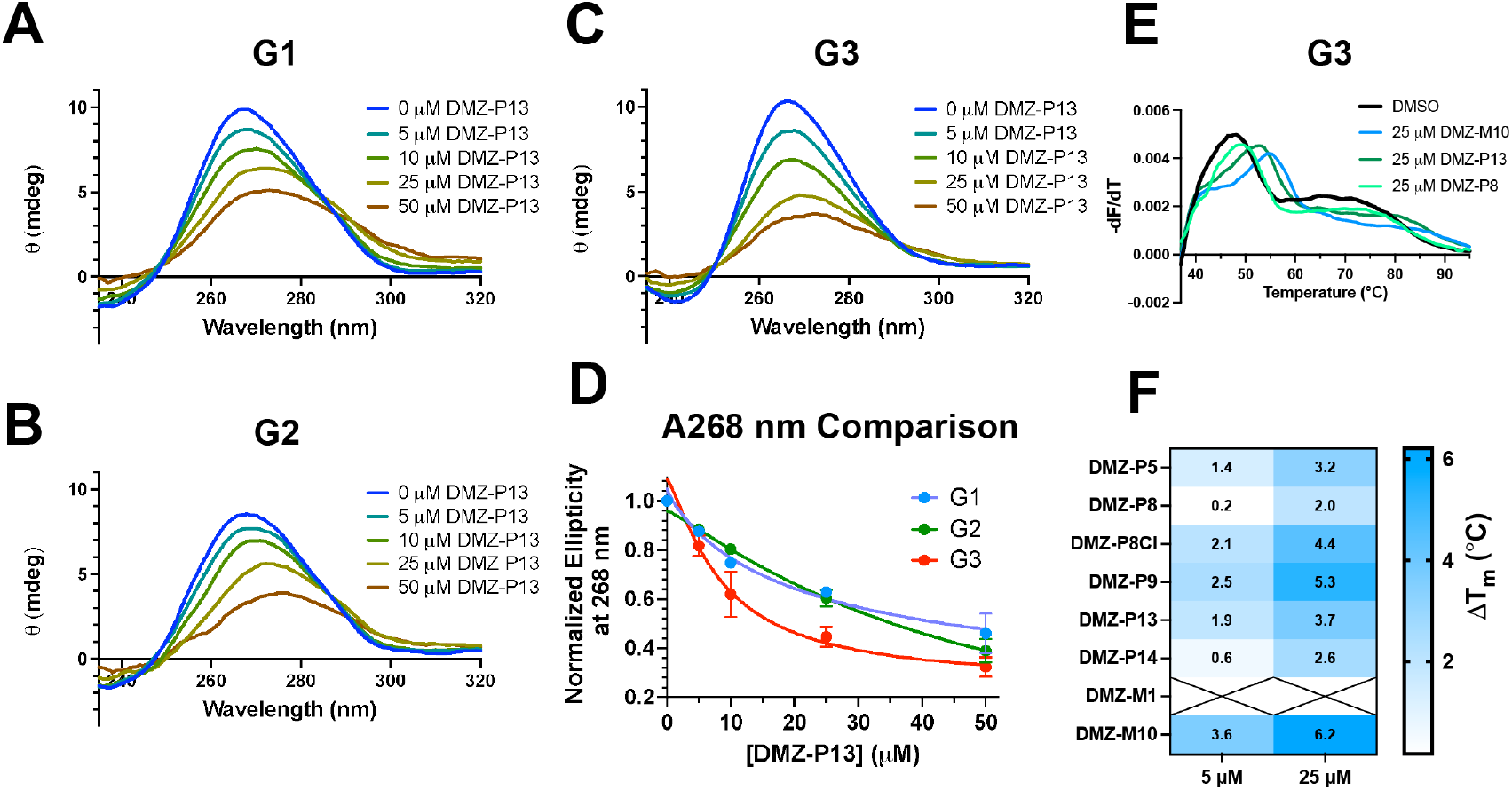
A-C) **DMZ-P13** titrations with 1 µM **G1, G2**, and **G3** independently. Experiments were carried out in biological duplicate and graphs are representative replicates. Buffer used was consistent with IDA (20 mM HEPES-KOH, 52 mM KCl, 0.1 mM MgCl_2_ pH 7.4). D) Non-linear curve-fits of the normalized ellipticity values at 268 nm for the three rGQs upon **DMZ-P13** addition demonstrates left shifting and approaching saturation for **G3** and not other MALAT1 rGQs, recapitulating the **DMZ-P13** selectivity. E) Representative **G3** DSF curves incubated with either DMSO control (*black*), **DMZ-M10** (*blue*), **DMZ-P13** (*green*), or **DMZ-P8** (*light green*). Experiments were performed in independent triplicate of three technical replicates. F) DMZ library hits for **G3** at 5 and 25 µM. DSF experiments were carried out in 20 mM HEPES-NaOH, 52 mM NaCl, 0.1 mM MgCl_2_ pH 7.4. Individual DSF curves are in **Figure S7**.

The combined IDA and CD data were consistent with **G3** being a MALAT1 rGQ with a higher propensity for small molecule targeting and modulation when compared to **G1** and **G2**. As all four, including the **N1** rGQ, retain the canonical three quartet rGQ structure, the most striking difference between these rGQs is the composition and length of their loop regions. **N1** has four one adenine loops, **G2** has two G-rich four nucleotide loops, **G1** has a five nucleotide A-rich loop, and **G3** has a seven-nucleotide pyrimidine-rich loop (**Figure 1**). The differences in small molecule binding suggests that the loop size plays a role in the binding mode of action.^33,64^ Interestingly, these differential interactions are further supported by previously reported molecular dynamic simulations, where DMZ molecules preferred binding to parallel GQs with longer loop regions.^64^ Beyond the high affinity for **G3**, the DMZ binding trends further corroborated this loop length trend, with the average CD_50_ increasing with decreasing loop length (**G1** < **G2** < **N1**). While DMZ molecules bind to **G3** preferentially over the other rGQs, each small molecule likely binds in a unique way based on the substituents, suggesting that a more elaborate study of binding mode would be beneficial in further elucidating how these DMZ molecules interact with rGQs and impact structure stability.

#### Differential Scanning Fluorimetry

To measure the melting temperatures of our **G3** construct, we used differential scanning fluorimetry (DSF).^70^ This approach has been used previously to assess the thermal stability shifts of DMZ molecules on the MALAT1 TH^59,60^ as well as small molecules on DNA GQs.^71^ For DSF evaluation, we initially tested an array of HEPES-based buffers with high K^+^, low K^+^, Li^+^, Na^+^, and Mg^2+^ for impacts on **G3** melting profiles. We observed shifts in melting temperature (T_m_) for **G3** consistent with previous rGQ metal cation studies (For T_m_, high K^+^ > low K^+^ > Li^+^ > Na^+^) (**Figure S6**).^9^ Magnesium quenched the fluorescence of RiboGreen™ and was not pursued further. We decided to test small molecule shifts on **G3** thermal stability in a sodium-based buffer, with the hypothesis that it would promote a larger dynamic range of observable thermal stability changes.

We used the previously identified DMZ hits from IDA to test for changes in thermal stability against **G3**. The DMZs demonstrated consistent dose-dependent stabilization and right shifting of the melting peak, as indicated by positive ΔT_m_ values (**Figure 3E-F, S7**). Furthermore, shifts in **G3** thermal stability were not correlated to affinity as measured by IDA, as **DMZ-M10** and **DMZ-P9** had the greatest increases but more moderate affinity when compared to **DMZ-P13** and **DMZ-P5**. Thermal stabilization did not match the conformational shifts suggested by CD, which showed reduced ellipticity upon **DMZ** addition, which is consistent with GQ destabilization based on literature reports.^49^

#### In vitro RT-qPCR

rGQs have been observed to inhibit readthrough by ribosomes *in cellulo* and are therefore regulated by helicases in many biological processes.^51,52,72,73^ Similarly, inhibition of polymerase read-through using reverse transcription-based methods has facilitated the identification of rGQs transcriptome-wide.^15,74,75^ Therefore, modulation in the rGQ structural stability could be linked to the ability of these enzymes to read through an rGQ forming sequence. Similar to the method developed by Useugi and co-workers for NRAS rGQs^32^ and more recently by our laboratory for the MALAT1 TH,^76^ *in vitro* RT-qPCR assays can be used to assess differences in *in vitro* folding of RNA structures as well as evaluate small molecule-driven structural modulation. DMZ binding that modulates the rGQ structure would be anticipated to impact the amount of cDNA produced by the reverse transcriptase (RT) when compared to a DMSO/vehicle control. For instance, an rGQ stabilizer would be expected to decrease the amount of cDNA produced (**Figure 4A**). By measuring the amount of cDNA by qPCR, a ΔC_t_ value can approximate the degree to which the small molecule stabilizes or destabilizes the structure of the rGQ. This ΔC_t_ value is derived from the cycle thresholds (C_t_), or the cycles at which the measured fluorescence signal of the SYBR dye crosses the threshold value, a value set over the background fluorescence of the dye (**Figure 4A**).

**Figure 4.**
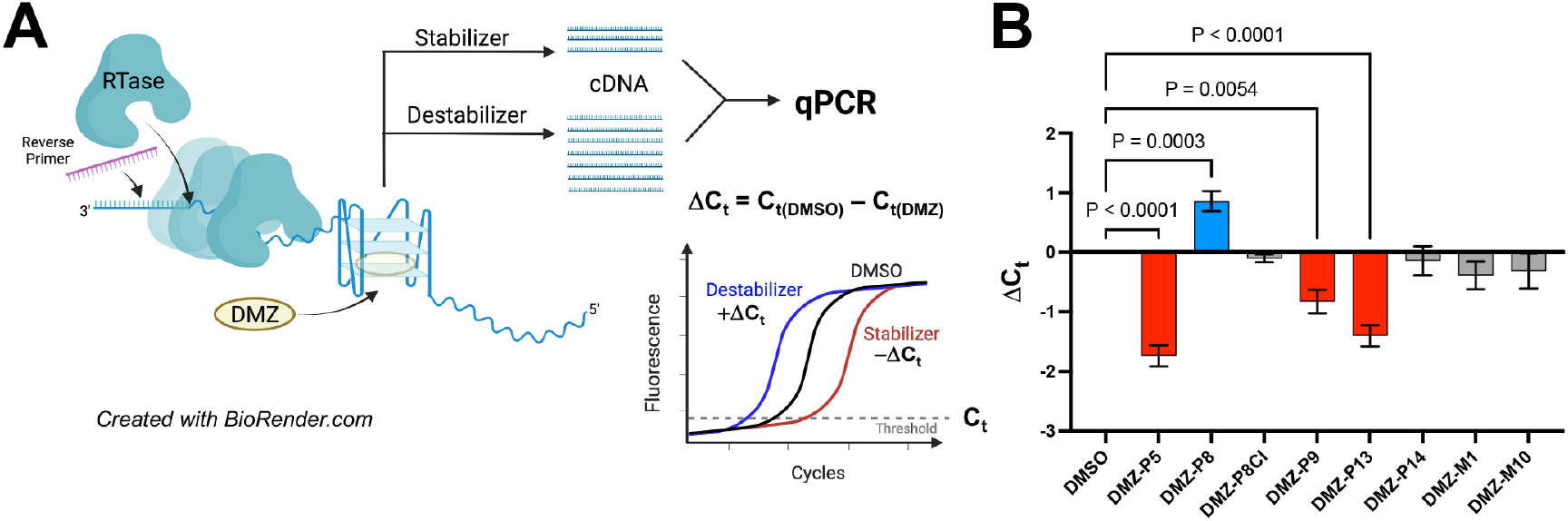
A) Schematic of RT-qPCR assay, created with BioRender.com. The stability of the rGQ structure directly decreases the ability of the RTase to read through the RNA transcript and produce cDNA, providing a readout for how the DMZ small molecules impact the structure. B) ΔC_t_ values with standard error of the mean (SEM) for **G3** with DMZ IDA hits, calculated from a DMSO control. One-way ANOVA was used to identify statistically significant changes in **G3** stabilization. Reverse transcriptase reaction conditions were: 2.5% DMSO, [**G3**] = 1 µM, [DMZ] = 5 µM, 10 units SSIV (ThermoFisher), 150 nM reverse primer, 600 µM dNTPs, 500 nM MgCl_2_. The qPCR conditions were: 4 µL RT reaction, 300 nM forward primer, 300 nM reverse primer, 10 µL SYBR Fast qPCR mix (KAPA), and nuclease-free water to 30 µL, plated in technical triplicate. Representative curves are in **Figure S8**.

We evaluated change in RT read-through for **G3** with the IDA hits. We used a longer **G3** construct for this method, increasing the flanking regions to limit competition between structural folding and primer binding. Interestingly, this approach revealed statistically significant **G3** stabilizers including **DMZ-P13, DMZ-P5**, and **DMZ-P9** and one ‘destabilizer’, **DMZ-P8** (**Figure 4B, S8**). Beyond those four, there were no significant changes in readthrough after small molecule addition. There was no correlation to affinity, as we see the highest stabilization for **DMZ-P5** rather than the expected order, **DMZ-P13** < **DMZ-P5** < **DMZ-P9** < **DMZ-P8**. The fact that **DMZ-P8** and **DMZ-P13** displayed opposite shifts suggested a difference in binding mode based on the additional ethylene spacer of **DMZ-P13**, which presumably increases flexibility for an ideal fit. **DMZ-P9** also has this ethylene linker, whereas **DMZ-P8** does not and may prefer loop binding and favor an alternative conformation more prone to read-through (**Figure 2B**). This trend is consistent with thermal stability, as **DMZ-P9, DMZ-P13**, and **DMZ-P5** demonstrated strong increases in ΔT_m_, whereas **DMZ-P8** had little to no change. These data further highlight the need to consider rGQ-small molecule interactions holistically to identify promising small molecule probes, as again affinity may not be linked to desired function.

### Stability-based proteomics evaluates rGQ-GQBP interactions

We adapted the workflow used in our previous study for measuring RNA impacts on protein stability for the rGQ and small molecule comparisons (**Figure 5A**).^77^ We started by collecting nuclear lysate samples from three unique LNCaP cell passages and prepared six different treatments: 1) **G3**, 2) **G3 + DMZ-P8**, 3) **G3 + DMZ-P13**, 4) **DMZ-P8**, and 5) **DMZ-P13**, and 6) lysate (**L**) control. We then applied a SPROX ‘one-pot’ protocol,^78^ allowing analysis of the average methionine peptide oxidation per treatment, giving a readout of domain-level protein stability.

**Figure 5.**
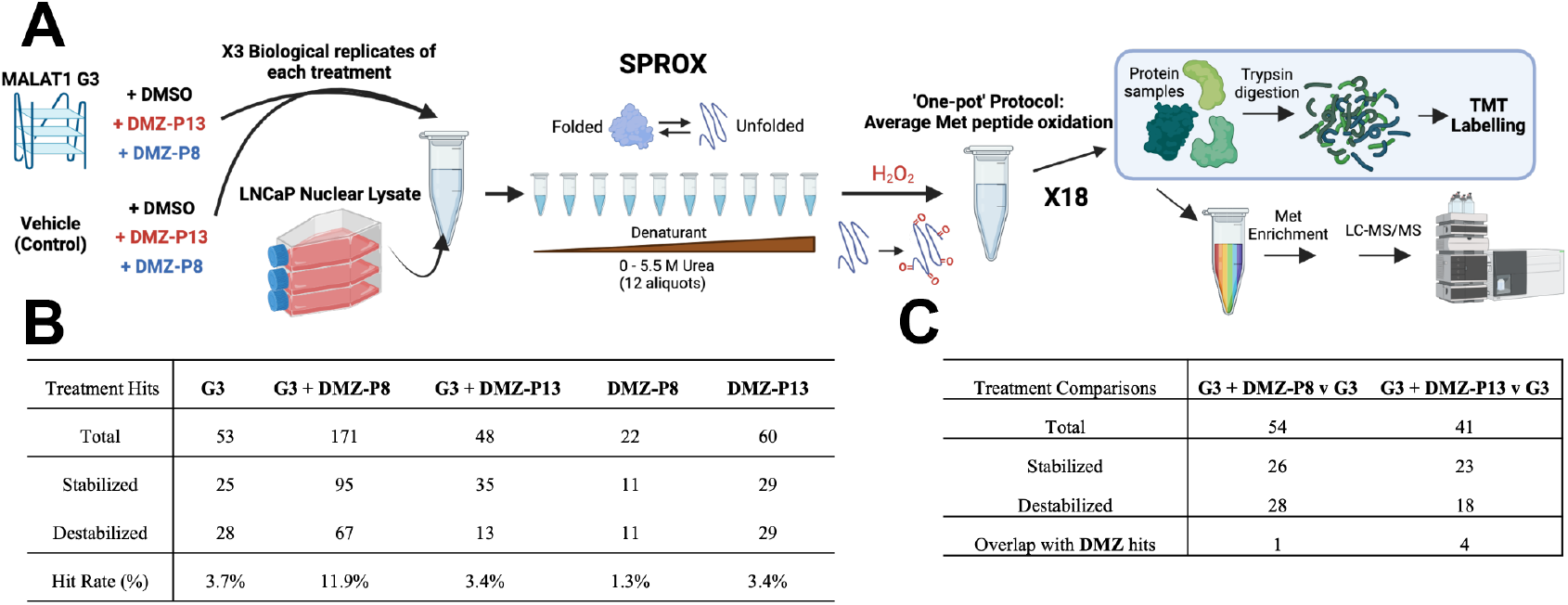
A) Workflow of the **G3** SPROX experiment, created with BioRender.com. B) All assayed **G3 +/-DMZ** protein hits versus lysate. Hits were defined as peptides having a Z-score greater than 2 (stabilized) or less than -2 (destabilized), as well as a p-value < 0.05. Hit rate (%) was calculated based on the total proteins assayed in Experiment **A** or **B** for the respective treatments. C) Evaluation of all assayed protein hits which demonstrated statistically significant changes in Z-score between **G3 + DMZ-P8/DMZ-P13** and the **G3** treatment. This identifies proteins that are impacted by small molecule addition. Overlap with DMZ hits refers to overlap within these comparisons to **DMZ-P8 v L** and **DMZ-P13 v L** hits. The overall lack of overlap suggests that the majority of hits being identified for statistically significant changes in stability are therefore **G3** + **DMZ** dependent. Hit rate (%) was calculated based on Experiment **A** protein numbers for each of these two comparisons. Stratifying these results by RNA-binding and GQBP binding can be found in **Table S4**.

Across these experiments, we assayed a total of 1927 unique proteins (Experiment **A** assayed 1432 proteins, Experiment **B** assayed 1753 proteins) and 8,888 unique methionine peptides (Experiment **A** assayed 5921 peptides, Experiment **B** assayed 7860 peptides) (**Table S1**). In line with previous work,^77^ we defined our hit criteria as peptides that had a Z-score greater than 2 standard deviations from the mean when compared to **L** (**v L**) and yielded a p-value < 0.05 in a student’s t-test.

Protein hit counts for each of the treatments when compared against the lysate-only control (i.e. **G3 v L** for **G3** hits) are shown in **Figure 5B**. Protein hit rates were calculated by dividing the number of hits per comparison by the total assayed in that specific experiment (**A** or **B**) (**Table S4**). These rates are comparable to those found from the previous study.^77^ We stratified these hits by GO-annotated RNA-binding functions^79,80^ and by known GQBPs collected by Zhou and co-workers^27^ and found that we identified hit peptides from high confidence RNA-binding proteins (RBPs) and GQBPs (**Table S4**).

Experiment **A** allowed for the comparison between **G3 + DMZ-P8**/**DMZ-P13 v G3** conditions, enabling the identification of hit peptides matching to proteins that shifted significantly in stability upon small molecule addition to the rGQ and lysate mixture. Focusing first on the distribution of hit peptide stability shifts in these comparisons, all assayed proteins reflected a near even split in stabilized and destabilized peptides, suggesting that general conformational changes were observed (**Figure 5C**). However, among RBPs, a shift was observed towards destabilized peptides, which is consistent these small molecule interactions leading to loss of GQ-GQBP interactions. To assess off-target small molecule effects, these proteins were analyzed for overlap with the hits from **DMZ-P8 v L** and **DMZ-P13 v L**. We saw only one RBP and GQBP overlapping hit in **G3 + DMZ-P13 v G3**, specifically AUF1, which is stabilized in the presence of **DMZ-P13**. However, this specific example was represented by different peptides in these two comparisons (**G3 + DMZ-P13 v G3** and **DMZ-P13 v L**) that have opposite stability shifts, suggesting that this rGQ and off-target small molecule interaction were through different modes of action. No overlapping hits were found for DMZ-P8. These results revealed that the majority of protein hits for **G3 + DMZ-P8/DMZ-P13 v G3** were likely not through off-target small molecule impacts.

*Characterization of known rGQ-protein interactions* SPROX identified 12 unique proteins annotated as RBPs and GQBPs that had significant changes in stability upon **G3** addition (**G3 v L**). These included proteins with well-characterized rGQ binding, including hnRNPF,^53^ hnRNPH1/3,^54,55^ EWSR1,^81,82^ TARDBP,^83^ and U2AF65 (**Table 1**, *bolded*).^47^ The identified hit peptides in these proteins were also identified within or in proximity to RNA binding domains in each of these proteins, as previously observed.^77^

**Table 1:** Identified RBP-GQBPs by SPROX mass spectrometry

| Accession Number | Protein Name | Gene Name | Number of G3 v L Peptide hits | Peptides identified | Stabilized/destabilized? | RNA binding domain? |
| --- | --- | --- | --- | --- | --- | --- |
| P17987 | T-complex protein 1 subunit alpha (TCP-1-alpha) (CCT-alpha) | TCP1 CCT1 CCTA | 1 | P17987 [154-159] | Destabilized (-2.03) | No |
| P26368 | Splicing factor U2AF 65 kDa subunit (U2 auxiliary factor 65 kDa subunit) (hU2AF(65)) (hU2AF65) (U2 snRNP auxiliary factor large subunit) | U2AF2 U2AF65 | 1 | P26368 [445-452] | Destabilized (-2.34) | RRM3 |
| P31942 | Heterogeneous nuclear ribonucleoprotein H3 (hnRNP H3) (Heterogeneous nuclear ribonucleoprotein 2H9) (hnRNP 2H9) | HNRNPH3 HNRPH3 | 1 | P31942 [233-261] | Stabilized (2.23) | RRM2 |
| P31943 | Heterogeneous nuclear ribonucleoprotein H (hnRNP H) [Cleaved into: Heterogeneous nuclear ribonucleoprotein H, N-terminally processed] | HNRNPH1 HNRPH HNRPH1 | 1 | P31943 [348-355]; P55795 [348-355] | Stabilized (3.14) | RRM3 |
| P41252 | Isoleucine--tRNA ligase, cytoplasmic (EC 6.1.1.5) (Isoleucyl-tRNA synthetase) (IRS) (IleRS) | IARS1 IARS | 3 | P41252 [419-428], P41252 [892-903], P41252 [911-917] | Destabilized (-2.79, -2.46, -2.92) | No |
| P48444 | Coatomer subunit delta (Archain) (Delta-coat protein) (Delta-COP) | ARCN1 COPD | 1 | P48444 [289-304] | Destabilized (-2.02) | MHD |
| P52597 | Heterogeneous nuclear ribonucleoprotein F (hnRNP F) (Nucleolin-like protein mcs94-1) [Cleaved into: Heterogeneous nuclear ribonucleoprotein F, N-terminally processed] | HNRNPF HNRPF | 1 | P52597 [91-98] | Destabilized (-3.25) | RRM1 |
| P54136 | Arginine--tRNA ligase, cytoplasmic (EC 6.1.1.19) (Arginyl-tRNA synthetase) (ArgRS) | RARS1 RARS | 1 | P54136-1 [72-79] | Destabilized (-2.14) | No |
| Q01844 | RNA-binding protein EWS (EWS oncogene) (Ewing sarcoma breakpoint region 1 protein) | EWSR1 EWS | 1 | Q01844 [269-292] | Stabilized (3.14) | RGG1 |
| Q12906 | Interleukin enhancer-binding factor 3 (Double-stranded RNA-binding protein 76) (DRBP76) (M-phase phosphoprotein 4) (MPP4) (Nuclear factor associated with dsRNA) (NFAR) (Nuclear factor of activated T-cells 90 kDa) (NF-AT-90) (Translational control protein 80) (TCP80) | ILF3 DRBF MPHOSPH4 NF90 | 2 | Q12906-7 [393-408], Q12906-7 [528-540] | Stabilized (2.63, 3.20) | DRBM1, DRBM2 |
| Q13148 | TAR DNA-binding protein 43 (TDP-43) | TARDBP TDP43 | 1 | Q13148-1 [84-95] | Stabilized (2.75) | RRM1 |
| Q15424 | Scaffold attachment factor B1 (SAF-B) (SAF-B1) (HSP27 estrogen response element-TATA box-binding protein) (HSP27 ERE-TATA-binding protein) | SAFB HAP HET SAFB1 | 1 | Q15424-1 [382-393] | Destabilized (-2.09) | RRM1 |

The Maiti and Kwok groups identified **G3** protein interactions through RNA-protein pulldowns including nucleolin, nucleophosmin, NONO, and DHX36, though there was no further characterization of the potential binding mechanism or functional impact.^49,50^ We did not observe nucleolin, nucleophosmin, or NONO as hits upon addition of **G3**, though these proteins did appear as hits in the **G3 + DMZ-P8** condition; however, similar Z-score shifts were seen with **DMZ-P8** alone for these three proteins. We believe that the lack of **G3** interaction with these three proteins in our experiments may result from our decision to not treat the lysates with micrococcal nuclease, which is typically done to remove endogenous RNA, in order to retain a competitive environment for our RNA substrates. As these proteins have high affinity for endogenous rGQs,^36,49,50,84^ **G3** may not have been able to compete for binding and therefore not impact the stability of these proteins over the lysate control. On the other hand, the small molecule could then bind to these endogenous RNAs and lead to the observed stability change such as in this case for these three proteins with **DMZ-P8**.

Beyond the filtering by RNA-binding functions and GQBPs, there were other proteins that warranted further investigation. One of these proteins included Histone 1.0, a protein involved in DNA binding and nucleosome organization.^85,86^ Interestingly, the peptide identified was from the N-terminal domain of Histone 1.0 and demonstrated the greatest Z-score stabilization of all **G3 v L** hits. This N-terminal domain is positively charged as well as intrinsically disordered and has been proposed to interact with components of the spliceosome. This stabilized peptide suggests one of two hypotheses: 1) this N-terminal domain can bind rGQs, leading to stabilization of the intrinsically disordered domain and protection of the methionine from oxidation, or 2) the N-terminal domain is interacting with other proteins with which **G3** also interacts, for example U2AF65, hnRNPF, hnRNPH, among others,^85^ which then leads to methionine stabilization through **G3**-protein scaffolding. In both cases, these hypotheses point to potentially novel roles for the N-terminal domain of Histone 1.0 to bind to rGQs, or splicing factors which coordinate with rGQs, to allow for its functional epigenetic regulation and additional functions.

Separately, a peptide from PurB transcription factor was also identified. This protein belongs to a family of evolutionarily ancient transcription factors that bind to purine-rich sequences, best characterized in c-Myc regulation.^87^ This peptide, found in a family-conserved Region 3 of this protein, has been previously determined to bind to DNA GQs. This SPROX peptide also maps to a known synthetic peptide that has been used to target C9orf72 repeats.^88^ With this finding, we went back through the data looking for another characterized synthetic GQ-targeting peptide derived from [53-105] in DHX36.^18,89^ We identified this N-terminal peptide of DHX36 as a hit in the **G3 + DMZ-P13 v L** comparison, but this peptide did not meet the hit criteria in **G3 v L**. Visualizing this peptide within the crystal structure (PDB: 5VHE) shows the methionine peptide (DHX36 [61-68]) directly interacting with the crystallized GQ.^89^ This example further demonstrates the ability of SPROX to identify peptides already known to target rGQs, suggesting that some other identified peptides, like the N-terminus peptide of H1.0 if hypothesis (1) holds true, could be used for potential rGQ targeting.

### Small molecules differentially modulate rGQ-GQBP interactions

We next conducted a detailed analysis of how DMZ molecules impacted **G3**-protein interactions on a global scale. When we compare the RNA-binding **G3 + DMZ-P8/DMZ-P13 v G3** hits to **G3** hits directly, we found three overlapping hits for the **DMZ-P8** comparison and one for **DMZ-P13** based on our statistical cutoffs, showing modulation of ∼15% of the observed **G3**:protein interactions (**Figure 6**, *right*). As changes in protein stability are related to RNA-protein affinity, **DMZ-P8** and **DMZ-P13** may not directly compete with some **G3**-protein interactions at the tested concentration (50 µM) as these can bind with low nanomolar affinity.^84^ While numbers are small, the greater overlap between **G3 + DMZ-P8 v G3** and **G3 v L** hit proteins is consistent with the destabilizing **DMZ-P8** leading to greater changes in rGQ structure, i.e. favoring alternative conformations that then lead to larger changes in **G3**-protein interactions relative to the stabilizer **DMZ-P13**. Notably, a small number of proteins became hits upon small molecule addition, *i*.*e*., were significantly stabilized only by the combination of **G3** and small molecule, and these are also discussed below.

**Figure 6.**
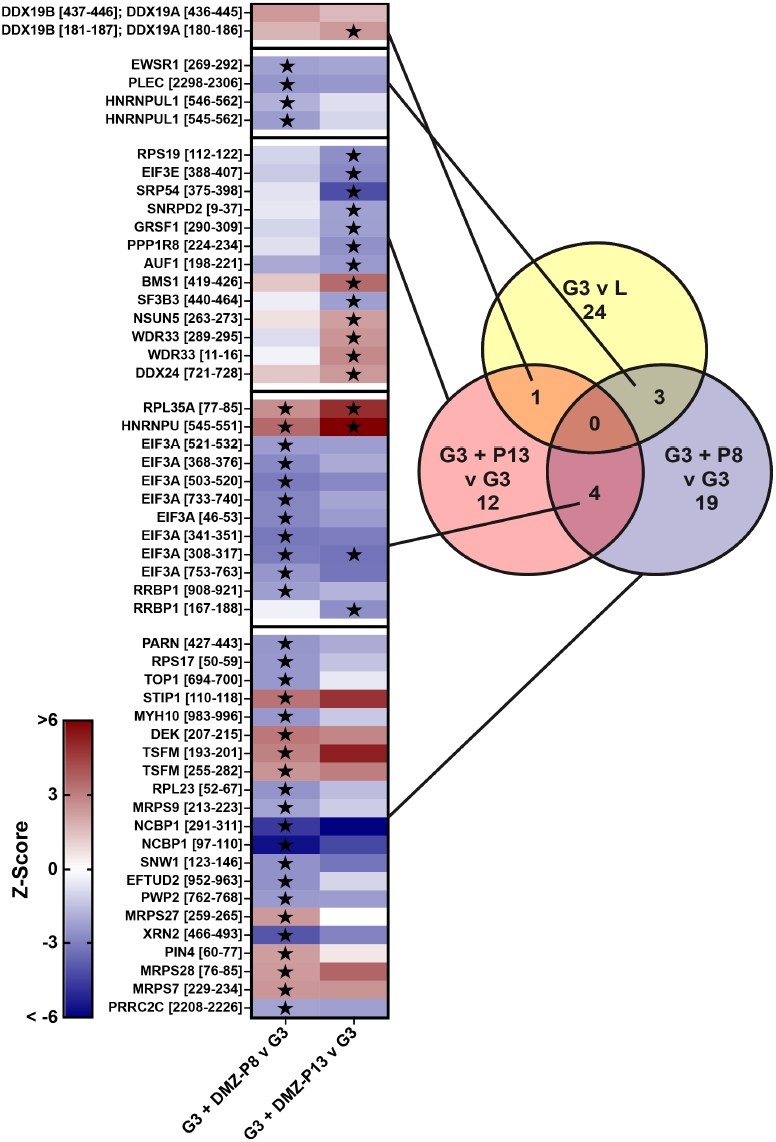
*Right*) Venn diagram of RNA-binding protein hits and overlap across **G3 v L** and **G3 + DMZ-P8/DMZ-P13 v G3** comparisons. Comprehensive lists of these proteins and corresponding peptides are available in **Table S2** and **Figure S10**. *Left*) Z-score comparisons for all peptides identified as hits found for **G3 + DMZ-P8/DMZ-P13 v G3** comparisons, with the locations of these peptides in the quandrants of the venn diagram drawn. Hits in each comparison are denoted by black stars.

#### DMZ-induced GQ-GQBP displacement

The four proteins that overlap between **G3 v L** and **G3 + DMZ-P8/DMZ-P13 v G3** include DDX19A/B, hnRNPUL1, PLEC and EWSR1 (**Figure 6**, *Left*). Interestingly, as the hit peptides in these proteins were evaluated for their respective Z-scores compared to lysate, we observed changes in Z-score likely linked to rGQ-GQBP displacement. To elaborate, each of these peptides demonstrated a > 2 or < -2 Z-score for **G3 v L**, as was necessary to be called as a hit in that comparison. However, upon DMZ addition, the Z-score moved towards zero, consistent with stability at lysate levels (identified through the respective **G3 + DMZ-P8/DMZ-P13 v G3** comparisons, noted by the peptide’s color). These shifts are consistent with the DMZ molecules inhibiting the rGQ-GQBP interaction. With DDX19A/B for example, the peptides map to a helicase ATP-binding domain ([125-295]) and the helicase C-terminal domain ([306-474]), respectively, suggesting displacement of the **G3**-induced ‘destabilization’ across both domains with DMZ addition. PLEC and EWSR1 demonstrate this displacement for both small molecules though the changes are identified as statistically significant in **G3 + DMZ-P8 v G3**. For all four proteins, the **DMZ-P8 v L** and **DMZ-P13 v L** did not demonstrate peptide stability shifts to the same degree as **G3 v L** alone, suggesting that these proteins were specifically modulated through small molecule-induced changes to **G3**-GQBP binding.

hnRNPUL1 is a particularly interesting hit as it has recently been associated with skeletal and developmental regulation and ALS.^90,91^ Strikingly, there is no annotated RNA-binding domain close to these SPROX peptides. Upon further investigation, we found that the two overlapping peptides identified here ([546-562], [545-562]) overlap with a recently characterized mutation site identified in humans and zebrafish to cause skeletal deformations.^90^ This mutation causes a frameshift that leads to an early stop codon, significantly shortening the protein in close proximity to the peptides we identify in our study. These peptides are also near the C-terminal end of a dead polynucleotide kinase domain, and hnRNPU-like proteins have an intrinsically disordered C-terminal domain which may also be contributing to binding, leading to these high positive stability shifts in **G3 v L**.^91^ However, these peptides demonstrate a **G3 + DMZ-P8 v L**-unique decrease in stability change, whereas **G3 + DMZ-P13 v L** retained the stability observed in **G3 v L** (**Figure S10**). The stability-shifts observed suggest small molecule-dependent destabilization, where **DMZ-P8**, the destabilizer, disrupts the **G3**-hnRNPUL1 interaction and **DMZ-P13** does not. Looking more closely at the data, the five total SPROX peptides from this protein (three of which are **G3 v L** hits, all in the region of [518-562]) recapitulate this Z-score trend (**Table S2**).

#### DMZ-induced GQ-GQBP stabilization

We also saw peptides that were not identified in **G3 v L** but **G3 + DMZ-P8** and/or **G3 + DMZ-P13** treatments changed their stability to the point they were selected as hits. This result leads to the hypothesis that these proteins may be shifting in stability through the formation of a ternary complex between the protein, **G3** and **DMZ-P8/DMZ-P13**. Examples are shown in **Figure 6C**, including peptides from known RNA-binding proteins and GQBPs, including hnRNPU and EIF3A. These two peptides are facilitated by both **DMZ-P13** and **DMZ-P8** addition to have a greater stability shift from lysate than **G3 v L** alone. We also identify specific DMZ-**G3**-protein interactions that reflect this interaction stabilization. For instance, GRSF1 and WDR33 show **DMZ-P13** specific changes in stability when **G3** is present. On the other hand, **G3 + DMZ-P8 v G3** identifies XRN2 and PARN as two proteins with peptides that shift significantly upon **DMZ-P8** addition, but not significantly when **DMZ-P13** is added.

#### DMZ off-target effects

As mentioned previously, we did not consider **DMZ-P8 v L** and **DMZ-P13 v L** hits in **G3 + DMZ-P8/DMZ-P13 v G3** comparisons due to likely interference of **DMZ-P8**/**DMZ-P13** off-target interactions. However, analysis of these small molecule hits reveal potentially valuable off-targets to consider for later cell-based uses of these DMZ molecules, and some of the strongest peptide stability shifts relative to lysate were observed for DMZ addition only (**Figure S9**). These hits included strong destabilization of peptides in Y-box binding protein 1 and POLRA1, proteins primarily involved in cellular stress responses and ribosomal RNA transcription.^92-94^

Through this SPROX dataset, we demonstrate that small molecule stabilizers and destabilizers can both similarly displace or stabilize the same rGQ-GQBP interactions or selectively favor and disfavor different interactions through small molecule-specific modulation of rGQ stability. Our work constitutes the first application of stability-based proteomics to assess RNA-small molecule-protein interactions in equilibrium, working to decipher how small molecules ultimately influence rGQ-GQBP recognition on the proteomic level and to inform future therapeutic avenues for targeting rGQ-GQBP interactions. This proteomic analysis is limited by the availability of methionine peptides in the desired interaction domains and interfaces; however, the number of peptides in or in proximity to RNA binding domains supports the use of this technique to identify potential GQ-GQBP interaction sites and facilitate their investigation. Pulling from the unique examples above such as with hnRNPUL1, this dataset and technique sets the groundwork to investigate GQ-GQBP interactions on the global scale, as well as provide stabilization-based evidence as to how small molecules can ultimately shift phenotypes based on changes in protein-binding profiles.

## CONCLUSIONS

Through holistically evaluating rGQ-DMZ interactions, then further applying those insights to rGQ-GQBP interactions on the proteomic level, we provide a workflow for comprehensively evaluating rGQ-targeting small molecule impacts on RNA-protein interactions. Our different biophysical datasets highlighted members of our DMZ library as novel binders for rGQs and identified **G3** as a MALAT1 rGQ with a higher propensity for RNA-small molecule targeting, likely due to increased loop length. Our data further support that the binding of DMZ hits **DMZ-P8** and **DMZ-P13** differentially destabilize and stabilize the **G3** fold, respectively, despite arising from the same scaffold. Through the use of stability-based proteomics, we investigated how these rGQ-folding modulators impact **G3**-protein interactions, providing both a new protein interactome for **G3** and hypotheses to understand how differential GQ-modulating ligands can foster or deter rGQ-GQBP interactions.

In addition to the specific interactions uncovered as small molecule targets such as **G3**:hnRNPUL1 and **G3**:EWSR1, these methods enable investigations into the nature of stabilizing vs. destabilizing rGQ ligands across a range of different rGQ structures and the global analysis of how these modulations differently influence rGQ:protein interactions, resulting in changed RNA function. We expect these insights to apply to targeting of other RNA tertiary structures and that these methods will significantly advance our understanding of specific mechanisms of action in small molecule-RNA targeting.

## EXPERIMENTAL SECTION

### *In vitro* transcription and sequences

DNA duplex template sequences shown below in **Table S5** were purchased with standard desalting or SDS PAGE purification from Integrated DNA Technologies (IDT). DNA duplex templates were initially diluted to 100 µM then further diluted to 50 µM. The sequence was then *in vitro* transcribed (IVT) using the following protocol. For a given 3 mL IVT reaction, the following were added in order: 300 µL of rNTP mix (25 mM of each, NEB) 18.75 µL MgCl_2_ (1 M), 120 µL Tris-HCl (1 M at pH 8.0), 75 µL of spermidine (0.1 M), 30 µL of Triton-X (0.1%), 30 µL of DTT (1 M), 600 µL of Q5 GC Enhancer (5x), 15 µL of pyrophosphatase enzyme (100 U/mL), 1.655 mL of RNase-free water, 6 µL of DNA duplex template (50 μM) and 150 µL of T7 polymerase enzyme kindly provided by the Tolbert lab at Case Western University. The 26bp-flanking **G3** had one difference, with Triton-X being replaced by 300 µL of DMSO (100%) and RNase-free water adjusting to an addition of 1.385 mL. The reaction was then incubated at 37 ^°^C for 2 hours. Following incubation, the reaction is treated once with 300 µL DNase I buffer (10x) and twice with 120 µL DNase I (New England Biolabs) in intervals of 30 minutes, followed by addition of 10% of the reaction volume of EDTA (0.5 M). The desired RNA is then extracted using phenol chloroform extraction (1:1 volume) and further purified via ethanol precipitation (added 2:1 volume of 100% molecular biology grade ethanol, placed at -80 °C for 30 minutes, then spun down at 4000 xg for 30 minutes at 4 °C). All RNA samples were lyophilized overnight and then resuspended in 150 µL RNase-free water, then RNA samples were further cleaned using the Zymo Research™ RNA Clean & Concentrator-100 kit. 12bp-flanking MALAT1 rGQs were used for IDA, WaterLOGSY, CD, DSF, ThT, and SPROX experiments. 26bp-flanking **G3** was used in RT-qPCR experiments. Purity and size of the RNA construct was confirmed by Invitrogen™ E-Gel Power Snap Electrophoresis System. Samples were heated to 95 °C for 5 minutes prior to gel loading, and 20 µL were loaded on the gel. 1% E-Gels to confirm purity were run for 10 minutes, stained with SYBR Safe DNA stain, and imaged on the same system. Oligo primer sequences were purchased from IDT and dissolved to an initial 100 µM stock for RT-qPCR experiments.

### Indicator displacement assay screening and titrations

This protocol was adapted from previous reports from our laboratory.^60,65^ First, rGQs were annealed at 95 °C for 5 minutes and on ice for 10 minutes. These rGQs were then serial diluted in HEPES buffer (20 mM HEPES-KOH, 52.6 mM KCl, 0.1 mM MgCl_2_ pH 7.4) in a 96-well plate in triplicate. 8 µL of each dilution was transferred to a 384-well plate followed by 8 µL of a 500 nM solution of RiboGreen™ dye (Invitrogen) in the same buffer. The plates were excited at 487 nm (8 nm slit) and emission was recorded at 530 nm (8 nm slit, focal height 11.3 mm) using a CLARIOstar plate reader (BMG labtech). The affinity of the dye for each rGQ was determined by fitting the raw fluorescence in GraphPad Prism (version 10.2.0 (335)) using the four-variable agonist versus response nonlinear curve fit function.

The determined affinity of the rGQ was used as the concentration of rGQ for DMZ screening, assuming 0.5 dye fraction bound (**G1**: 0.446 µL, **G2**: 0.397 µL, **G3**: 0.0326 µL, **N1**: 0.674 µL). Five concentrations (1, 5, 10, 25, 50 µM) of all DMZ molecules were used for initial screening against all four rGQs, again performed in HEPES buffer (20 mM HEPES-KOH, 52.6 mM KCl, 0.1 mM MgCl_2_ pH 7.4) with 500 nM RiboGreen™. rGQs were allowed to anneal at their EC_50_ concentrations at 95 °C for 5 minutes and on ice for ten minutes, followed by incubation with 500 nM RiboGreen™ for 20 minutes. Screening was performed by transferring small molecules using an Echo 550 Acoustic liquid handler (Labcyte) in a 384-well plate. Then, 10 µL of rGQ at the concentrations noted above and 500 nM of RiboGreen^TM^ was dispensed in the 384-well plates using a ThermoFisher liquid dispenser. After centrifuging the plates at 4,000 rpm for 1 minute, the plate was allowed to incubate in the dark for 20 minutes. The plates were excited at 487 nm (8 nm slit) and emission was recorded at 530 nm (8 nm slit, focal height 11.3 mm) using a CLARIOstar plate reader (BMG labtech). Percent fluorescence indicator displacement (%FID) was calculated by subtracting and, subsequently, dividing by the blank wells with RNA-dye complex and no small molecule as shown in equation below.

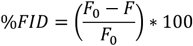

Hits were defined as those molecules which demonstrated dose-dependence increases in % fluorescent indicator displacement for one or more rGQs. Follow-up small molecule titrations were performed similar as to described above, except with 10-point titrations (0, 1.25, 2.5, 3.75, 5, 7.5, 10, 25, 37.5, 50, 75 µM) for the eight DMZ hit molecules selected. After carrying out the protocol again as described above, %FID values were plotted using GraphPad Prism (version 10.2.0 (335)), and CD_50_ values were determined by using the four-variable agonist versus response nonlinear curve fit function. These titrations were performed as a duplicate of technical triplicates, and CD_50_ values were determined from an average of two independent experiments.

### WaterLOGSY

Protocol was developed with input from previously published procedures.^67,68,95,96^ 8 µM 12-bp **G3** RNA was diluted from a lyophilized powder stock in NMR buffer (50 mM d11-Tris-HCl, 50 mM KCl pH 6.7 in 90:10 RNase-free water:D_2_O) for 300 µL per experiment. The G3 sample was heated at 95 °C for 5 minutes and slow cooled to room temperature for 1.5 hours. A reference 1D-1H and 1D WaterLOGSY spectrum were collected for **DMZ-P13** at a concentration of 1 mM in NMR buffer or 5% DMSO in NMR buffer. WaterLOGSY experiments were run by selecting the pulse sequence ephogyno.2 and manually adjusting the FID area to minimize the water signal. Upon successful collection of the reference spectra (16 scans), a second solution was made by titrating small molecule in 270 µL of 8 µM **G3** solution, adding 10 µL of small molecule solution every 5 minutes, to reach a final volume of 300 µL at a concentration of 7.2 µM of **G3** and 1 mM of **DMZ-P13**. A second WaterLOGSY experiment was then run (16 scans, 0.5-1.5s mixing time) and the spectra with and without RNA were overlapped. The reference peak of clear water was present to facilitate phasing. The current protocols have been optimized on a Bruker AVANCE III 700Hz at 25 °C. All data were visualized and processed with TopSpin instrument software (v. 8.2).

### Circular dichroism: rGQ validation and small molecule titrations

CD spectra were collected for initial validation and for small molecule titration using an Aviv model 62DS circular dichroism spectrometer. The rGQs were prepared and annealed for CD by snap-cooling, this entailing being heated at 95 °C for 10 minutes, then placed on ice for 30 minutes, and at ambient temperature for 45 minutes. 3.2 mL of 1 µM rGQ were prepared in this protocol, with 3 mL added to a quartz cuvette (path length: 1 cm). Initial validation studies were set at 1 µM strand concentration and were annealed in 10 mM Tris-HCl pH 7.4 with additions noted when assessing metal dose-dependence. Samples were scanned from 320 nm to 200 nm, 1 nm per scan for 1 s in triplicate. Buffer signal was subtracted from each sample before plotting using GraphPad Prism software (version 10.2.0 (335)). Data was smoothed using the smooth function (5 neighbors, second derivative).

For small molecule titrations, rGQ samples were diluted to 1 µM strand concentration and were annealed in 20 mM HEPES-KOH, 52 mM KCl, 0.1 mM MgCl_2_ pH 7.4. Small molecule stocks were initially diluted to 5 mM concentrations in 100% DMSO and were titrated into the samples to reach the desired concentrations. Control titrations with DMSO alone were taken to show that DMSO did not significantly impact the spectra (data not shown). At 50 µM small molecule, the total percent DMSO was 1%. 1% dilution of the original RNA stock was also taken into consideration. Each of the spectra shown are from two independent replicates performed on different days. Small molecule affinity was determined through normalization of the 268 nm ellipticity maximum (mdeg) to 0 µM molecule added and plotting these normalized values against small molecule concentration. Spectra smoothing and non-linear curve-fitting were again performed using GraphPad Prism (version 10.2.0 (335)).

### ThT fluorescence assay

ThT fluorescence experiments were performed on a CLARIOstar plate reader (BMG Labtech). rGQ samples were first annealed in 10 mM Tris-HCl, 40 mM KCl pH 7.4 using a slow-cooling protocol. On an Eppendorf Mastercycler Nexus™ Thermocycler, 500 mL of 4 µM rGQ samples were initially heated for 10 minutes at 95 °C, then let to cool at a rate of 1 °C/minute until it reached room temperature. Samples were then removed and 1 µL of 1 mM ThT (dissolved in 10 mM Tris-HCl pH 7.4) were added to the samples. 150 µL of this sample was added in triplicate to wells on a Greiner™ black round-bottom 96-well plate and was allowed to incubate and rotate in the dark at room temperature for 30 minutes. After incubation, the samples were placed on a centrifuge to spin down for 30 seconds at 4,000 rpm, then placed on the plate-reader. Samples were excited at 420 nm (8 nm slit) and emission was recorded at 490 nm (8 nm slit, focal height of 11.3 nm). Plates were gain adjusted prior to reading, setting the highest fluorescent well to 80% of the maximum reading. ThT ratios were determined by averaging the wells containing only a ThT control and using this value as F_0_. The rGQ samples were averaged by the technical triplicate and this value (F_rGQ_) was divided by F_0_ to leave the ratios. Ratios were the product of two independent replicates of technical triplicates for each MALAT1 rGQ.

### Differential scanning fluorimetry assay

Differential scanning fluorimetry experiments were performed as previously reported by our laboratory.^59,60^ All experiments were performed in white 96 well plates in a LightCycler® 96 (Roche). In a typical experiment, 4 µM of rGQ was annealed in 20 mM HEPES-KOH, 52 mM NaCl, 0.1 mM MgCl_2_ pH 7.4 by heating at 95 °C for 5 minutes, snap-cooling on ice for 10 minutes, and leaving the solution at room temperature for 20 minutes. Then, 42 µL of this solution was aliquoted to wells and 5 µL of DMZ were added in technical triplicates (2.5 µL of water, 2 µL of DMSO, 0.5 µL of a 500 µM or 100 µM DMZ stock to get 25 and 5 µM respectively). DMSO control conditions were prepared with 2.5 µL of water, 2.5 µL of 100% DMSO. The mixture was left to incubate at room temperature for 5 minutes, after which 3 µL of a 20 µM RiboGreen™ (Invitrogen) dye stock solution in appropriate buffer was added to each well. This led to final concentrations of 3.36 µM rGQ, 25 and 5 µM DMZ, 1.2 µM RiboGreen™, 5% DMSO in 20 mM HEPES-KOH, 52 mM NaCl, 0.1 mM MgCl_2_ pH 7.4. The 96-well plate was then sealed with an optically clear foil and centrifuged for 1 minute at 4000 rpm prior to being placed in the instrument. The light cycler program was created by selecting the melting curve method; fluorescence intensity was monitored using the SYBR Green I/HRM dye filter combination (465-510 nm) from 37 – 98 °C at a ramp rate of 0.01 ºC/second with 150 acquisitions per ºC. Melting profiles were obtained by T_m_ analysis in the LightCycler® 96 software (SW 1.1).

The initial buffer DSF tests used the above protocol but used different HEPES-based buffers, specifically 20 mM HEPES-KOH, 52 mM KCl, 0.1 mM MgCl_2_ pH 7.4 (K^+^ buffer), 20 mM HEPES-KOH, 150 mM KCl, 0.1 mM MgCl_2_ pH 7.4 (high K^+^ buffer), 20 mM HEPES-KOH, 52 mM LiCl, 0.1 mM MgCl_2_ pH 7.4 (Li^+^ buffer), 20 mM HEPES-KOH, 52 mM NaCl, 0.1 mM MgCl_2_ pH 7.4 (Na^+^ buffer), and 20 mM HEPES-KOH, 52 mM KCl, 25 mM MgCl_2_ pH 7.4 (Mg^2+^ buffer). 4 µM of each rGQ was annealed in the desired buffer, Then, 42 µL of this solution was aliquoted to wells and 5 µL of 50% DMSO was added in technical triplicates. The rest of the protocol was performed as described above and led to the selection of the Na^+^ buffer for small molecule screening.

### *In vitro* RT-qPCR

RT-qPCR assay protocol was adapted from Uesugi and co-workers^32^ as well as from our laboratory.^76^ All experiments were performed in white 96-well plates in a LightCycler® 96 (Roche). We used a two-step RT-qPCR approach, starting with preparation of the reverse transcription reaction which then we added to a qPCR second step. First, we annealed enough volume for all reactions (19.3 µL x # of reactions) of 1 µM 26-nt **G3** with 150 nM of **G3** reverse primer in RNase-free water at 95 ºC for 5 minutes followed by 10 minutes on ice. Pulling from a 200 µM stock of each DMZ molecule (100% DMSO), we added 1 µL to 19.3 µL of annealed **G3** + reverse primer to reach a final concentration of 5 µM DMZ in 20.3 µL per tube in a PCR strip. This was allowed to incubate at room temperature for 15 minutes. A second master mix was prepared with 10 U of SuperScript IV (Thermofisher Scientific), 600 mM dNTPs (New England Biolabs), and 7.5 mM MgCl_2_. This was diluted with RNase-free water to (19.7 µL x # of reactions) and vortexed quickly. This second master mix was added to each reaction over an ice block. After again vortexing the samples and spinning down using a tabletop centrifuge, these samples were then placed in an Eppendorf Nexus™ thermocycler programmed to incubate the samples at 42 ºC for 30 minutes, followed by a heated enzyme inactivation at 98 ºC for 3 minutes, and finally a hold at 4 ºC.

The second step qPCR mix was prepared by making a master mix of 300 nM of 26-nt **G3** forward primer, 300 nM of 26-nt **G3** reverse primer, 1/3 v/v SYBR qPCR Mix (KAPA), and RNase-free water to 26 µL per replicate of each reaction, so 3x was needed for technical plate triplicates. Reverse transcription reaction was removed from the thermocycler, heavily vortexed, and then 4 µL of the reaction was added by multichannel to the qPCR mix over an ice block. The 96-well lightcycler plate was sealed with an optically clear foil and centrifuged for 30 seconds at 4000 rpm prior to being placed on the LightCycler^®^ 96 instrument. The lightcycler program was as follows in **Table 2**.

**Table 2:**
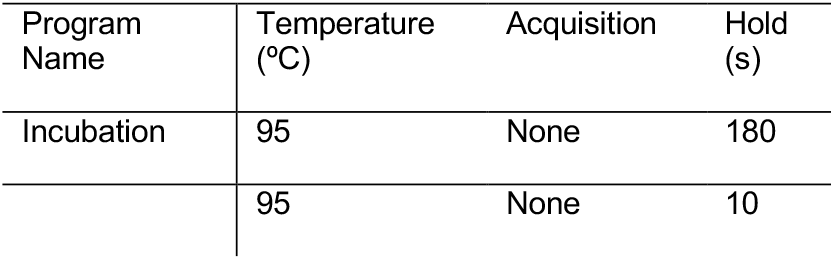
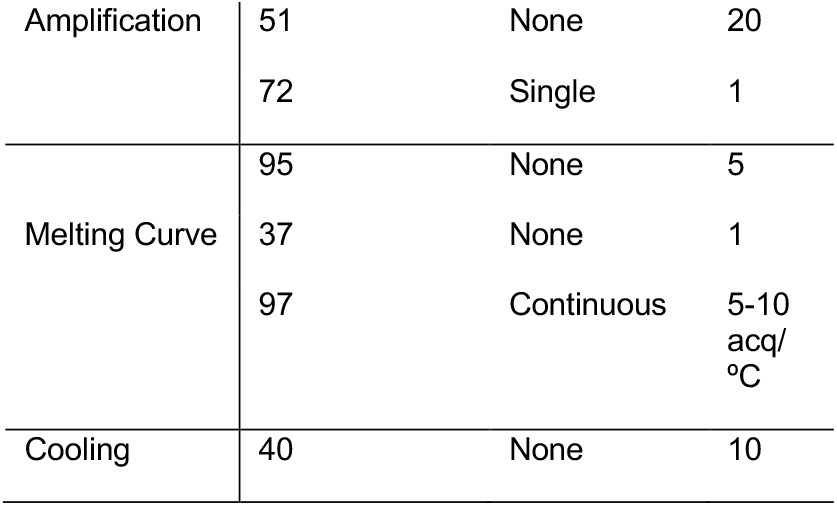
G3 RT-qPCR lightcycler program

Raw data were analyzed using LightCycler® 96 software (SW 1.1) using the absolute quantification analysis. C_t_ values were calculated using the software which sets a threshold value at 0.2 RFU. C_t_ values for DMSO were taken within each experiment and used as a reference point in order to calculate ΔC_t_ values for each DMZ molecule using the follow equation:

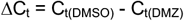

ΔC_t_’s were collected in at least a triplicate of technical triplicates across two or more days. A one-way ANOVA test was used to evaluate statistical significance against the DMSO control for each of the DMZ molecules tested using GraphPad Prism (version 10.2.0 (335)).

### Cell culture and nuclear lysate extraction

Cell preparation and SPROX methods were adapted from previous work.^77^ LNCaP prostate cancer cells were acquired from the Duke University Cell Culture Facility (originally from ATCC), STR-authenticated, and regularly checked for mycoplasma. Cells were maintained in Gibco RPMI 1640 with 10% Fetal Bovine Serum (FBS) without antibiotic or anti-mycotic until passage fifteen. Cells were incubated at 5% CO_2_ in a humidified incubator at 37 ^°^C. Cells were plated at a seeding density of approximately 3 x 10^6^ cells per T-175 flask. Cells were pelleted by first washing with PBS, then using 2 mL of Trypsin-EDTA evenly spread across the plate and allowing it to incubate for 5 minutes at 37 ^°^C. Cells were then removed from the plate with the addition of 10 mL of media, and cells were spun down at 180 rcf for 8 minutes. Media was disposed of, and cells were resuspended in 1 mL of PBS. These were then transferred to 1.5 mL Eppendorf tubes, spun at 10,000 rpm for 1 minute, and supernatant was pulled off and disposed. Cells were then frozen and stored at -20 ^°^C until use. Cell counts of cell pellets were approximately 20 x 10^6^ cells per pellet.

This nuclear lysate extraction protocol was adapted from “Nuclear Protein Extraction Without the Use of Detergent protocol” from Millipore Sigma.^97^ Cell pellets were gently resuspended in 1 mL of lysis buffer (10 mM HEPES-KOH, 10 mM KCl, 1.5 mM MgCl_2_ pH 7.9 supplemented with 1 mM DTT and 1X protease inhibitor cocktail (10X: 100 mM AEBSF, 2 mM leupeptin, 1 mM pepstatin A, 1.5 mM E-64, and 5 mM bestatin)) using a pipette. Suspended cells were allowed to sit on ice in lysis buffer for 15 minutes, before the cells were centrifuged at room temperature for 5 minutes at 450 rcf. The supernatant was removed, and the cells were resuspended in 400 µL of lysis buffer. A syringe with a narrow gauge (No. 25) hypodermic needle was used to draw the cell suspension into the syringe and then ejected with a single rapid stroke. This was repeated 6 times. These disrupted cells were centrifuged for 20 minutes at 11,000 rcf, and supernatant containing cytoplasmic lysate was removed. Nuclei were resuspended in 150 µL of Extraction buffer (20 mM HEPES-KOH, 420 mM NaCl, 1.5 mM MgCl_2_, 0.2 mM EDTA, 1 mM DTT, 1X protease inhibitor cocktail) with vortexing and the suspension was rotated for 30 minutes at room temperature. The suspension was centrifuged for 5 minutes at 21,000 rcf and supernatant was removed containing the nuclear lysate. The concentration was determined using the Qubit™ Protein Broad Range Assay (ThermoFisher). One LNCaP cell pellet provided 170 µL of approximately 3 mg/mL nuclear protein for downstream experiments.

### SPROX - Mass spectrometry: Sample preparation, data acquisition and analysis

12-bp **G3** RNA was first annealed in 150 mM KCl using 5-minute heat treatment at 95 ^°^C, followed by cooling on ice for 10 minutes. The SPROX treatments were prepared in a total volume of 130 µL and included 60 U of RNase Inhibitor (Invitrogen), 1% DMSO, 14.1 µM RNA (or equivalent volume of 150 mM KCl, vehicle control), 150 µM **DMZ-P8**/**DMZ-P13** for the small molecule treated samples (diluted from 1.5 mM stocks, 100% DMSO), and 3 mg/mL LNCaP nuclear lysate and made to volume with phosphate buffered saline (Corning). All six treatments were 1) lysate (**L**) only (DMSO + vehicle), 2) **G3** (DMSO + **G3**), 3) **G3 + DMZ-P8** (**DMZ-P8** + **G3**), 4) **G3 + DMZ-P13** (**DMZ-P13** + **G3**), 5) **DMZ-P8** (**DMZ-P8** + vehicle) and 6) **DMZ-P13** (**DMZ-P13** + vehicle). After treatments were prepared in biological triplicate, the samples were allowed to rotate at 4 ^°^C for 1.5 hours prior to running the SPROX protocol.

These treated samples were evaluated in SPROX experiments performed in biological triplicate using a one-pot SPROX method.^78^ In short, for each treatment replicate 360 µg of total protein was equally distributed across a series of 12 buffers containing 20 mM phosphate (pH 7.4), 150 mM NaCl, and final urea denaturant concentrations equally spaced in 0.5 M intervals from 0 to 5.5 M. The aliquoted treatments were diluted three-fold in the urea buffers and equilibrated for 2 hours at room temperature, at a final equilibration sample concentration of 4.7 µM **G3**, 20 U of RNase inhibitor, 0.33% DMSO, 50 µM **DMZ-P8**/**DMZ-P13**, 1 mg/mL LNCaP nuclear lysate, 20 mM phosphate pH 7.4, 150 mM NaCl, and desired urea concentrations. A methionine oxidation reaction was performed in each denaturant-containing sample through the addition of 3 µL of 30% (v/v)

H_2_O_2_. The final protein and H_2_O_2_ concentration in each denaturant-containing tube was 1 mg/mL and 0.9 M, respectively. The methionine oxidation reaction was allowed to proceed for 3 minutes before it was quenched with 250 µL of 1 M TCEP. Equal aliquots of the denaturant-containing protein samples in the denaturant series were combined in one tube. This was done for each of the 6 treatment conditions as well as 3 biological replicates for each treatment (6 total replicates of lysate control were prepared for two mass spectrometry experiments). The resulting 12 (treatments 1-4 for Experiment **A**) and 9 (treatments 1, 5-6 for Experiment **B**) protein samples were then prepared for LC-MS/MS quantitative bottom-up proteomic analysis using an isobaric mass tagging strategy involving 12 and 9 channels from a TMTpro 16-plex for Experiments **A** and **B** (respectively) and a previously described iFASP protocol.^98^ The methionine-containing peptides in the samples were enriched using a Pi^3^ methionine reagent kit following the manufacturer’s protocol (Nest Group). A C18 Macrospin column was used to desalt the labeled protein sample, which was ultimately subjected to triplicate LC-MS/MS analyses.

LC-MS/MS analyses of the samples generated in the SPROX experiments were performed on a Thermo Orbitrap Exploris mass spectrometer 480 system interfaced with a ThermoEasy nano LC 1200. A total of 1 µg of total peptide dissolved in 0.1% trifluoroacetic acid, and 2% acetonitrile in HPLC grade water to give a final concentration of 1 mg/mL was injected onto the column and subjected to an LC-MS/MS analysis using a nanoViper 2Pk C18, 3 µm, 100 Å, 75 µm X 25 cm analytical column (ThermoFisher) and gradient elution using a 90-minute linear gradient of 4 – 35% acetonitrile at a flow rate of 400 nL/min. The analytical column was maintained at a temperature of 45 ^°^C. A data-dependent acquisition was set up using a cycle time of 2.5 s. The resolution of the MS1 and MS2 data were at 120k and 45k, respectively. A normalized AGC target of 300% was used for both MS1 and MS2 analyses. The collision energy was set to 36% with an MS1 scan range of 375 – 1500 m/z, and an isolation window of 1.2 m/z with a dynamic exclusion duration of 45 s. Each peptide sample was analyzed in triplicate by LC-MS/MS (three biological replicates run in triplicate for each experiment).

The raw data from the LC-MS/MS experiments were searched against the 2017-10-25 release of the Uniprot Knowledgebase mammalian proteome using Proteome Discoverer 2.3 (ThermoFisher) and the SEQUEST HT database search algorithm. The TMT reporter ion intensities generated in each treatment replicate were normalized using a normalization factor generated from the summed reporter ion intensities in each individual replicate. Specifically, for SPROX, the non-methionine-containing peptide reporter ion intensities were used to normalize the methionine containing peptides. This step helps mitigate TMT channel to channel errors (i.e. differential sample loss and/or isobaric mass tag labelling efficiency).

As described previously,^77^ a ratio was generated for each treatment replicate and termed a fold-change value. Each of the treatment biological replicates were individually divided by the same biological replicate of the lysate/vehicle control to generate fold change ratios (e.g., F_avg(**G3**)_/F_avg(lysate)_ for **G3** versus lysate). These fold-change values were normalized for expression across the different biological replicates using the non-methionine containing peptide signals in a non-enriched sample, to allow for protein expression changes across the diversified cell passages. The fold-change values for each individual RNA treatment replicate were log_2_ base transformed, averaged, and subjected to a two-tailed Student’s t-test to calculate a p-value for each wild-type methionine containing peptide in SPROX. The hit peptides in SPROX were selected using a p-value less than 0.05 and a Z-score greater than 2 standard deviations away from the mean. Ultimately, due to the number of samples, this required two TMT experiments, termed **A** and **B**. Experiment **A** included three lysate controls, three **G3**, three **G3 + DMZ-P8**, three **G3 + DMZ-P13**. Experiment **B** included three separate lysate controls, three **DMZ-P8**, and three **DMZ-P13** alone treatments.

## ASSOCIATED CONTENT

### Data Availability

The mass spectrometry data generated in this work have been deposited to the ProteomeXchange Consortium via the PRIDE^99^ partner repository with the dataset identifier PXD060264.

### Supporting Information

The Supporting Information is available free of charge online. This includes:

Figures S1-S9, Tables S3-S4, and the synthetic procedure for the **DMZ-P8Cl** (PDF)

Summary of all assayed proteins in Experiments **A** and **B** in SPROX-MS (XLSX)

Summary of protein hit overlaps in Experiment **A** for the **G3 + DMZ-P8/DMZ-P13 v G3** comparisons (XLSX)

## AUTHOR INFORMATION

## Corresponding Authors

*Correspondence can be directed to Michael C. Fitzgerald; and Amanda E. Hargrove

## Present Addresses

^†^Department of Chemistry, University of Toronto, Mississauga, ON L5L1C6

## Author Contributions

The manuscript was written through the following contributions of all authors. J.G.M.: conceptualization, formal analysis, investigation, methodology, project administration, validation, visualization, writing (original draft, review and editing). M.Z.: conceptualization, formal analysis, investigation, methodology, writing (review and editing). M.A.B.: formal analysis, investigation, methodology, validation, writing (review and editing). M.D.Z.: investigation, writing (review and editing). N.I.M. and D.M.: investigation, methodology, validation. M.C.F.: supervision, funding acquisition, methodology, writing (review and editing). A.E.H: conceptualization, supervision, funding acquisition, methodology, writing (review and editing). All authors have given approval to the final version of the manuscript.

## Notes

The authors declare no competing financial interest.

## ACKNOWLEDGMENT

The authors thank Hargrove Laboratory members for their helpful conversations and support, including Drs. E. G. S. Hay and J. P. Falese for their feedback throughout project development. TOC graphic and Figures 1A, 4A, and 5A were created with BioRender.com.

## Table of Contents artwork

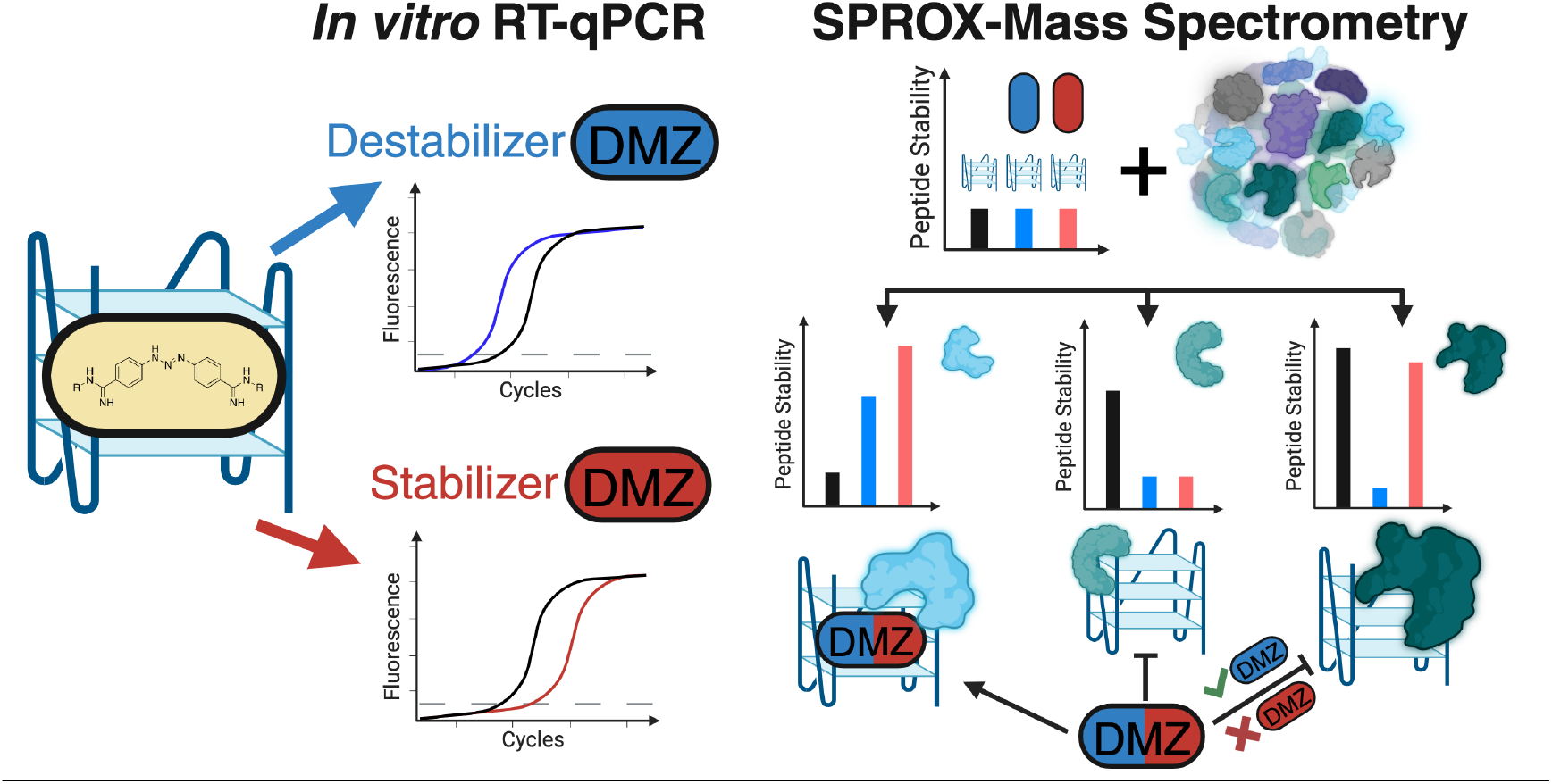

