## Supporting Information for "Small molecules reveal differential shifts in stability and protein binding for G-quadruplex RNA"

**Table of Contents:**


| | |
|---|---|
| **Figure S1. Circular dichroism validation** | **S-2** |
| **Figure S2. ThT validation** | **S-3** |
| **Figure S3. IDA curves** | **S-3** |
| **Figure S4. WaterLOGSY full figure** | **S-5** |
| **Figure S5. DMZ-P5 and DMZ-P13 CD titrations** | **S-6** |
| **Figure S6. DSF buffer studies for G1, G2, G3, and N1** | **S-7** |
| **Figure S7. DSF curves for DMZ hits against G3** | **S-8** |
| **Figure S8. Representative RT-qPCR curves** | **S-9** |
| **Figure S9. Z-score bar graphs for DMZ-P8 and DMZ-P13 off-target effects** | **S-9** |
| **Figure S10. Z-score heatmap for all G3 v L and DMZ+G3 v G3 peptides** | **S-10** |
| **Table S1: SPROX-MS Data** | **.xlsx** |
| **Table S2: Overlap between G3 v L and G3 + DMZ-P8/DMZ-P13 v G3** | **.xlsx** |
| **Table S3: DMZ structures, assay data and error** | **S-11** |
| **Table S4: Stratifying SPROX-MS hits by RNA-binding proteins and GQBPs** | **S-12** |
| **Table S5: Sequences and constructs used in this study** | **S-13** |
| **Synthesis of DMZ-P8Cl** | **S-14** |

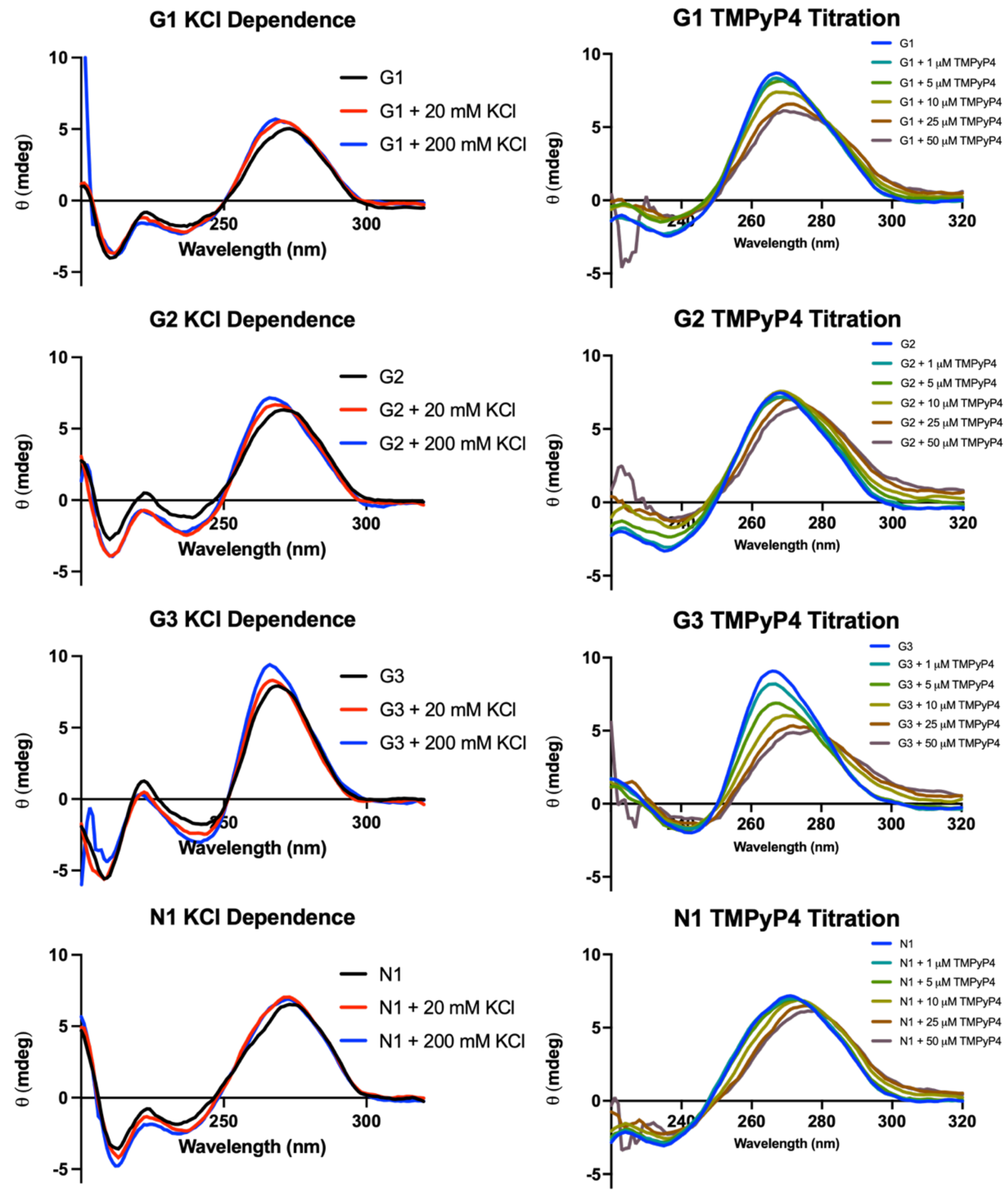


**Figure S1. rGQ CD spectra for KCl and TMPyP4 addition.** *Left)* Representative KCl titration for all rGQs. CD spectra was collected in 10 mM Tris-HCl pH 7.4 with KCl added to the concentrations noted, and KCl titrations were carried out in biological duplicate. *Right*) TMPyP4 titrations were run in 10 mM Tris-HCl, 200 mM KCl pH 7.4, and run in biological duplicate. Expected G-quadruplex spectra was recapitulated with the peak around 270 nm and trough around 240 nm.

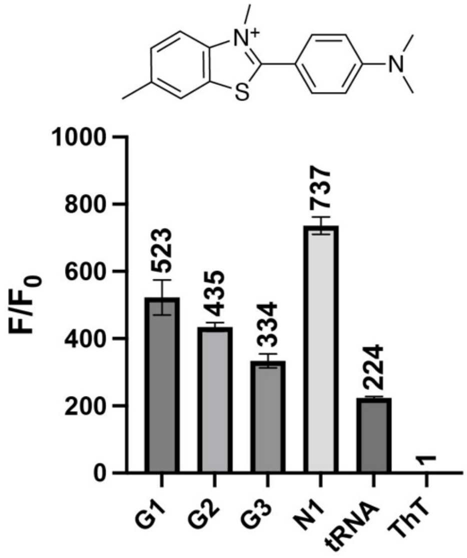


**Figure S2. ThT fluorescence assay shows increase with rGQs.** *Top*) Thioflavin T structure, *Bottom*) 4 μM **G1**, **G2**, **G3**, **N1**, as well as yeast tRNA were incubated with 2 μM ThT in 10 mM Tris-HCl, 200 mM KCl pH 7.4. F/$F_0$ was calculated by taking the raw fluorescence of **G1**, **G2**, **G3**, **N1**, and tRNA samples and dividing by ThT alone. Experiment was performed in biological duplicate, technical triplicate. The tRNA fluorescence change was comparable as a negative control as to what has been shown previously.[1]

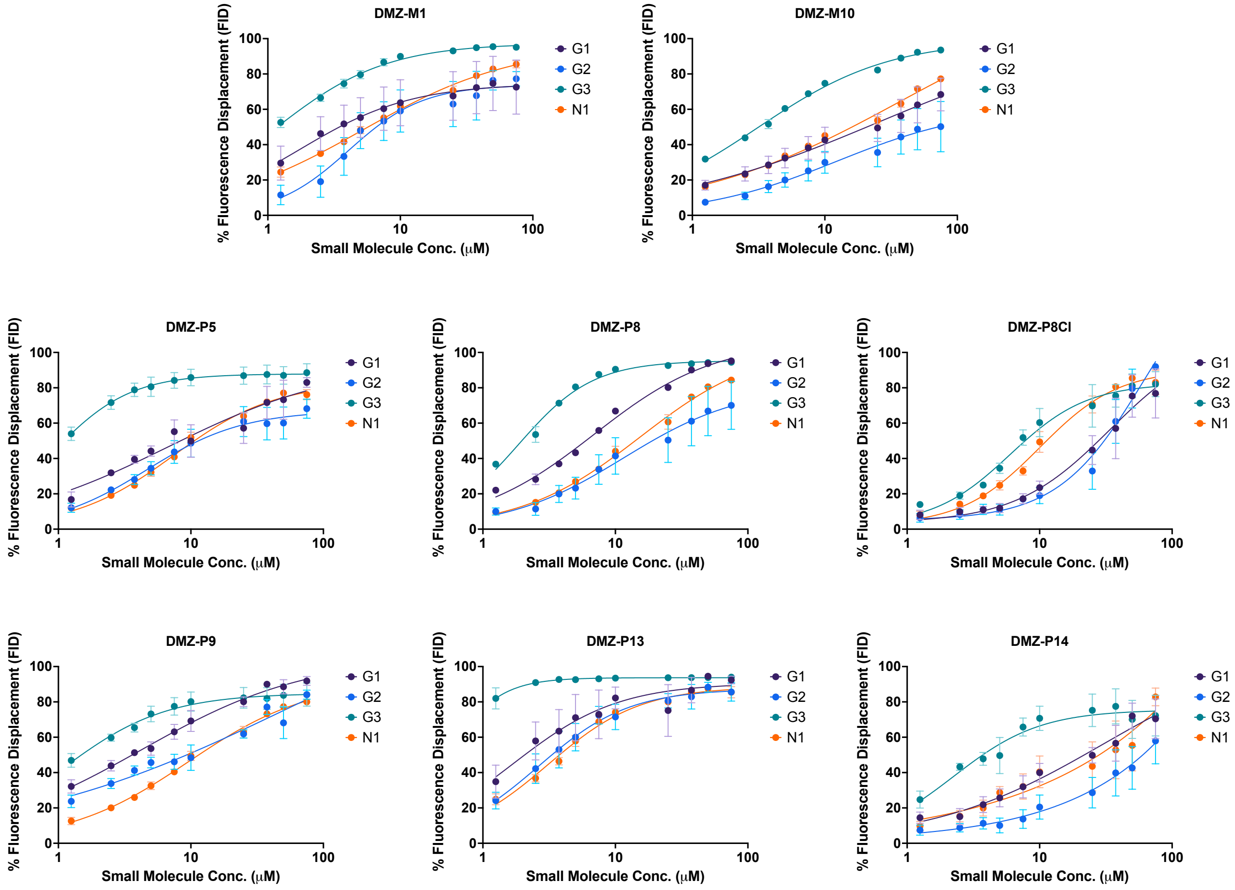

**Figure S3. IDA curves used to inform $CD_{50}$ values.** Each plot is the average of a triplicate of technical duplicates. DMZ concentration ranged from 1.25 - 75 μM. Average $CD_{50}$ values were determined through non-linear curve fitting each technical duplicate using GraphPad Prism (version 10.2.0 (335)) and then reporting the average and standard error of the mean of each experiment by selecting $CD_{50}$ values informed with a fit greater than $R^2$ = 0.8. These values are reported in **Table S3**.

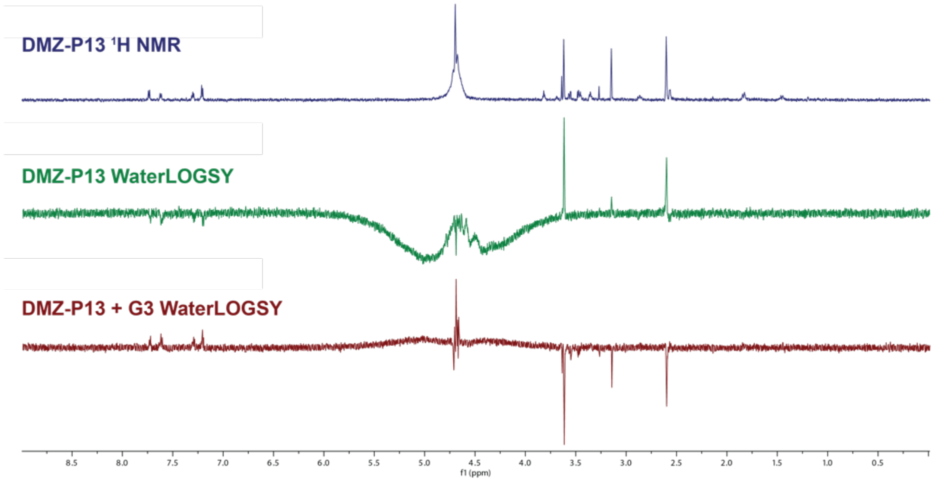
DMZ-P13 1H NMR
DMZ-P13 WaterLOGSY
DMZ-P13 + G3 WaterLOGSY
8.5 8.0 7.5 7.0 6.5 6.0 5.5 5.0 4.5 4.0 3.5 3.0 2.5 2.0 1.5 1.0 0.5
f1 (ppm)

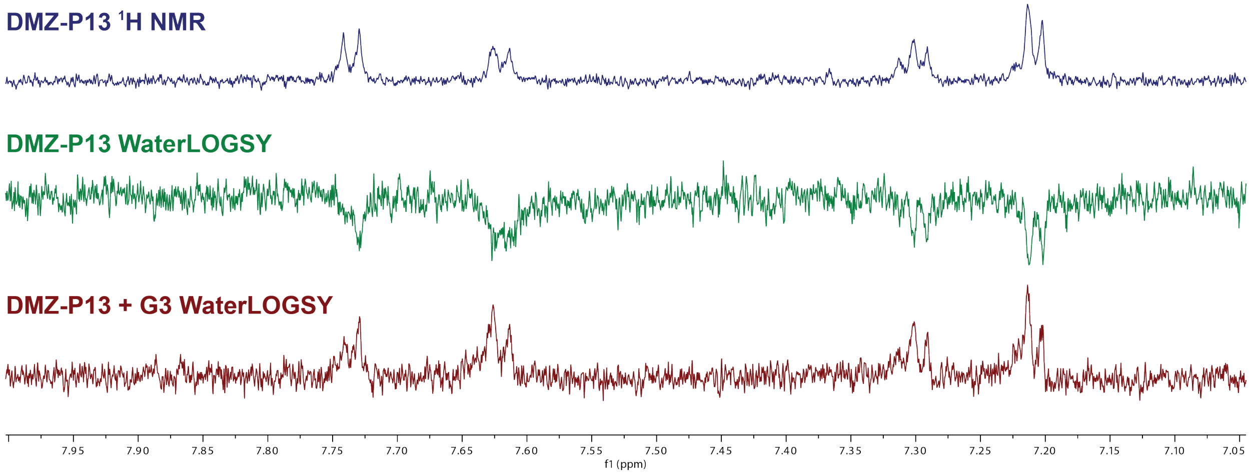
DMZ-P13 1H NMR
DMZ-P13 WaterLOGSY
DMZ-P13 + G3 WaterLOGSY
7.95 7.90 7.85 7.80 7.75 7.70 7.65 7.60 7.55 7.50 7.45 7.40 7.35 7.30 7.25 7.20 7.15 7.10 7.05
f1 (ppm)

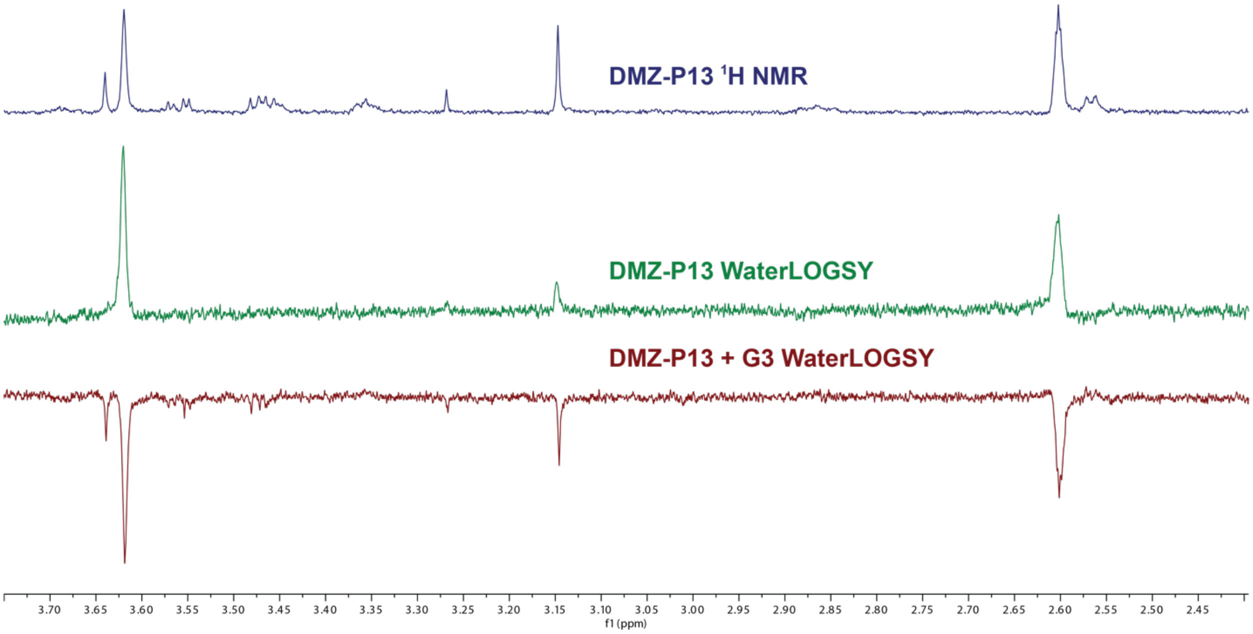
DMZ-P13 1H NMR
DMZ-P13 WaterLOGSY
DMZ-P13 + G3 WaterLOGSY
3.70 3.65 3.60 3.55 3.50 3.45 3.40 3.35 3.30 3.25 3.20 3.15 3.10 3.05 3.00 2.95 2.90 2.85 2.80 2.75 2.70 2.65 2.60 2.55 2.50 2.45
f1 (ppm)

**Figure S4. WaterLOGSY Spectra.** Spectra was collected as described with 7.2 μM **G3** and 1 mM **DMZ-P13**. In each figure, the blue spectra corresponds to the reference 1D-1H NMR, the green spectra is the 1D WaterLOGSY spectrum, and the red spectra corresponds to the combined 7.2 μM **G3** and 1 mM **DMZ-P13**. *Top*) The full spectra showing the in-phase water peak (~4.6 ppm) to control for peak inversion, *middle*) demonstrates a zoomed in region of the aromatic region showing the representative changes in peak inversion upon RNA addition, and *bottom*) shows a zoomed in look of the aliphatic region of the spectra.

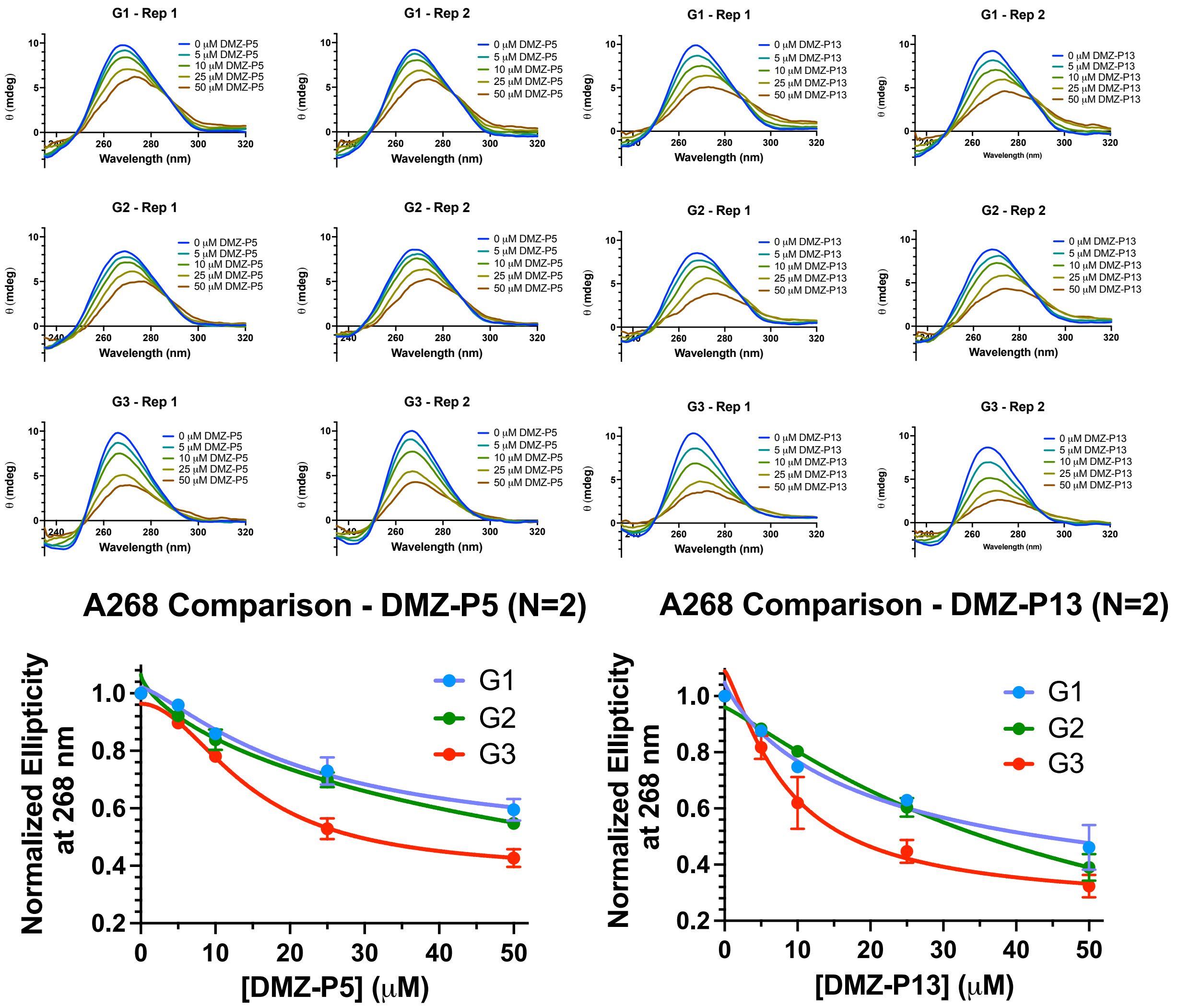


**Figure S5. CD spectra for DMZ-P5 and DMZ-P13.** Circular dichroism (CD) experiments were carried out in independent duplicates (Rep 1 and Rep 2) for each **G1**, **G2,** and **G3** at 1 μM. Raw CD signal was first background subtracted using a buffer blank, as well as confirmed that small molecule alone at the noted concentrations did not impact the spectra in the range of the canonical G-quadruplex spectra (data not included). Data was then smoothed in GraphPad Prism (version 10.2.0 (335)) using the smooth function (5 neighbors, second derivative) then plotted by small molecule concentration. **DMZ-P5** addition is shown on the left and **DMZ-P13** is shown on the right. *Bottom*) The absorbances at 268 nm was normalized to the 268 nm absorbance at 0 μM, then plotted the values based on concentration, producing the curves shown. These results recapitulated the affinity trends observed in the indicator displacement assay experiments, with a preference for **G3** binding over other RNA G-quadruplexes.

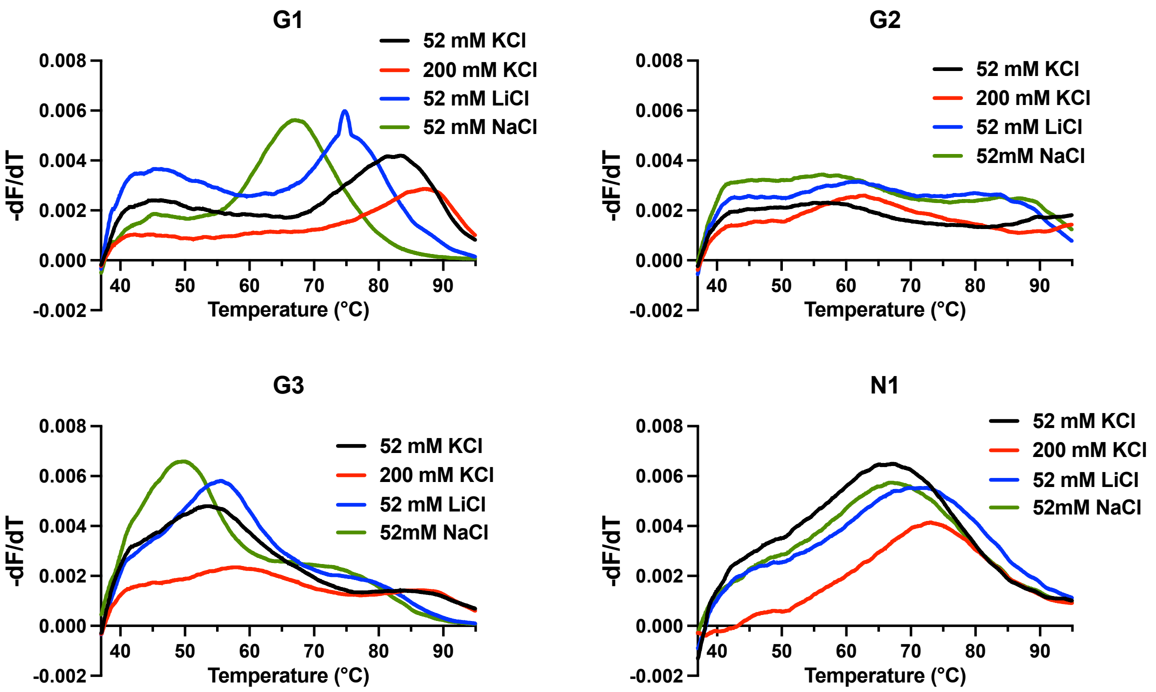


**Figure S6. DSF buffer evaluation of G1, G2, G3, N1.** Experiments were run in technical duplicate, and representative curves from each RNA G-quadruplex (4 μM) is shown. In the plots above, the derivative of the collected fluorescence over temperature was plotted against the temperature, highlighting the melting peaks. Black lines corresponded to the melts in 20 mM HEPES-KOH, 52 mM KCl, 0.1 mM $MgCl_2$ pH 7.4 ($K^+$ buffer), red lines corresponded to the melts in 20 mM HEPES-KOH, 150 mM KCl, 0.1 mM $MgCl_2$ pH 7.4 (high $K^+$ buffer), blue lines corresponded to 20 mM HEPES-KOH, 52 mM LiCl, 0.1 mM $MgCl_2$ pH 7.4 ($Li^+$ buffer), and green lines were melts in 20 mM HEPES-KOH, 52 mM NaCl, 0.1 mM $MgCl_2$ pH 7.4 ($Na^+$ buffer).

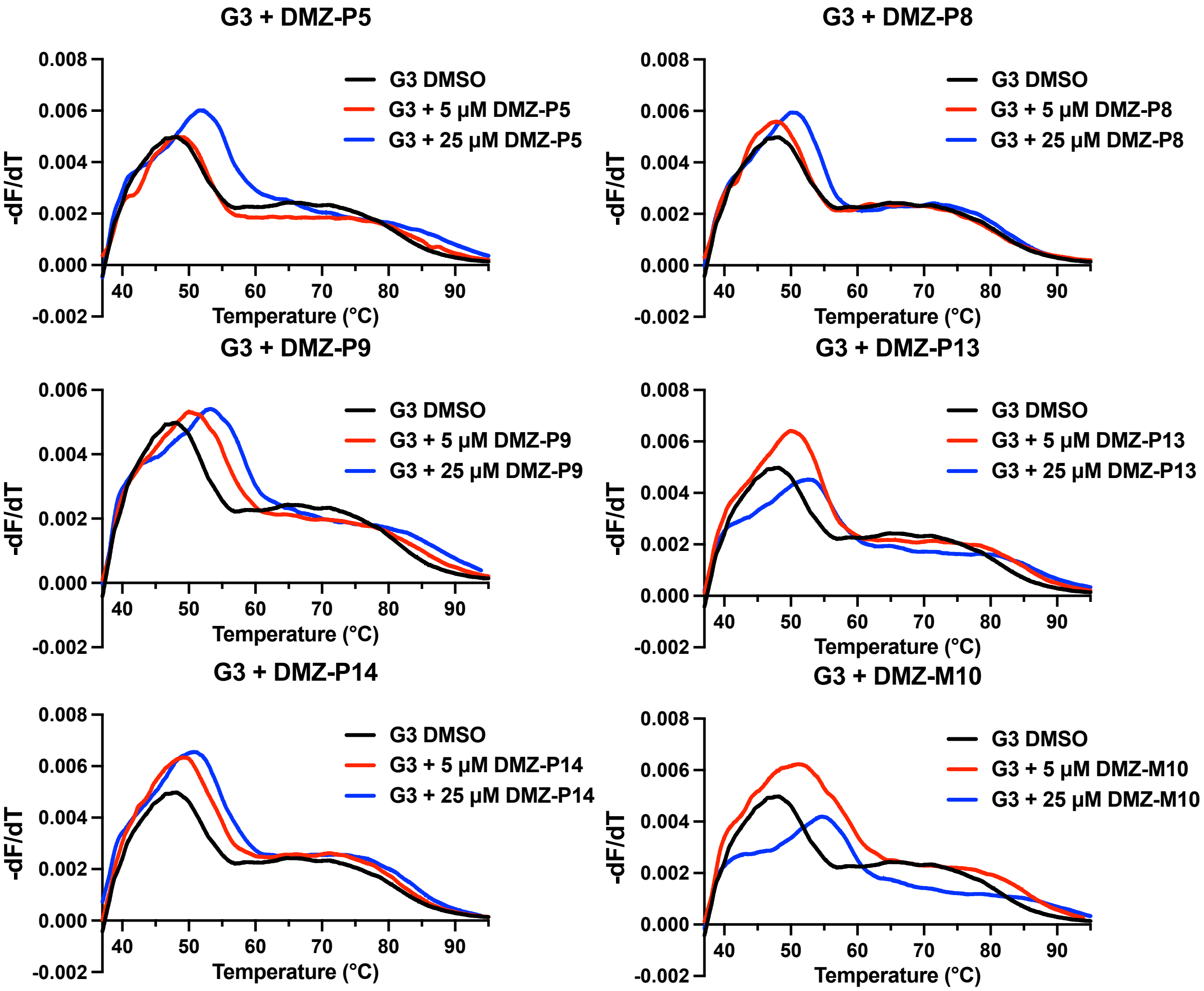


**Figure S7. DSF curves for DMZ hits against G3.** Plots are representative runs for six of the DMZ molecules noted from the DSF data for having dose-dependent interactions, as $T_m$ values were collected in biological duplicate of technical duplicate. In the plots above, the derivative of the measured fluorescence over temperature was plotted against the temperature, highlighting the melting peaks. Black lines corresponded to the melts in DMSO, red to 5 μM DMZ, and blue to 25 μM DMZ. Melts were carried out in 20 mM HEPES-KOH, 52 mM NaCl, 0.1 mM $MgCl_2$ pH 7.4 ($Na^+$ buffer) with 4 μM and 5% DMSO.

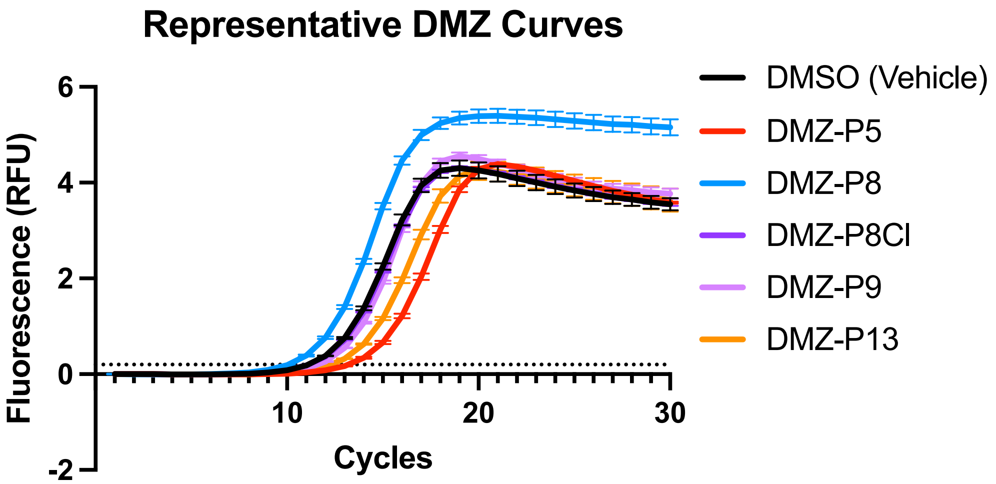


**Figure S8. Representative RT-qPCR curves.** Representative qPCR amplification curve of all significant molecules (and DMZ-P8Cl) informed by a single experiment with technical triplicate. Error bars shown represent standard deviation at each cycle. $C_t$ values were measured at a RFU threshold value of 0.2 RFU, shown here by the dotted line. $\Delta C_t$ values were calculated by taking the DMSO values from that experiment and subtracting the DMZ $C_t$ value from it, leading to stabilizers having (-) $\Delta C_t$ and destabilizers have (+) $\Delta C_t$. Experiments were run with an initial RT of 5 μM DMZ, 1 μM **G3**, 150 nM **G3** reverse primer, 2.5% DMSO, 600 mM dNTPs, 7.5 mM $MgCl_2$, and 10 U of SuperScript IV. This composition was allowed to proceed at 42 °C for 30 minutes, followed by 98 °C heat inactivation for three minutes. qPCR proceeded with 300 nM of forward and reverse primers, 1/3 (v/v) SYBR Fast qPCR mix, and 4 μL of the RT reaction. $\Delta C_t$ were informed by at least triplicate of technical triplicates.

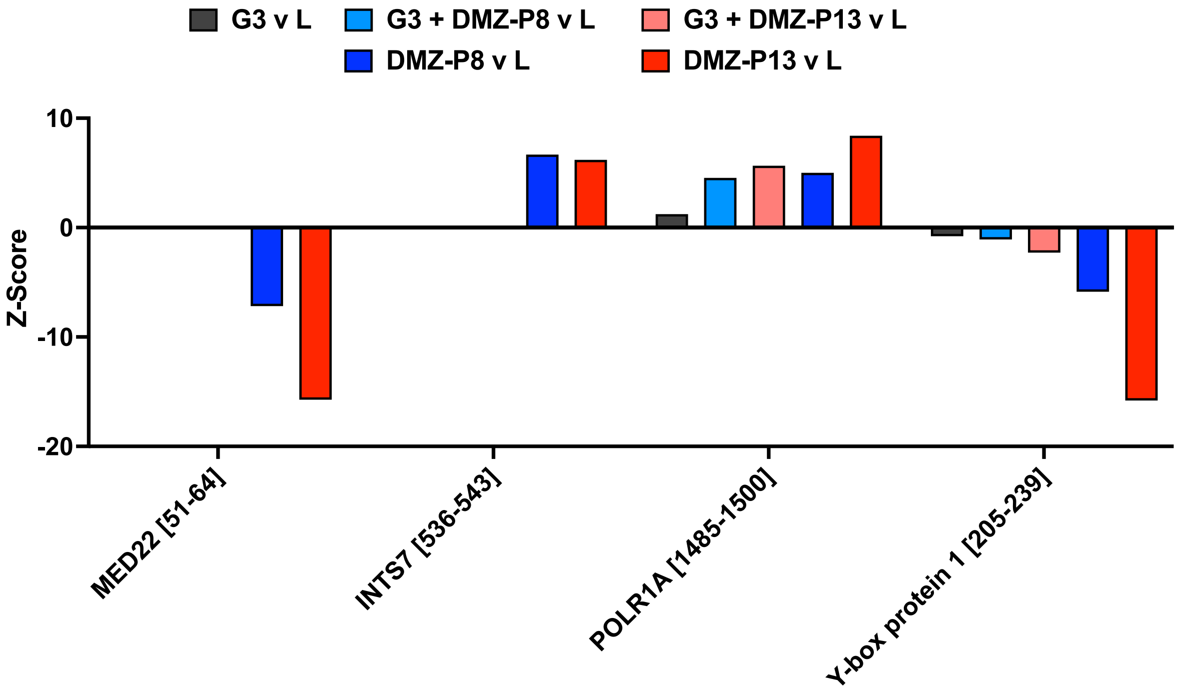


**Figure S9. Z-Score bar graphs for DMZ-P8 and DMZ-P13 off-target effects.** Z-Scores shown for select significant peptide hits for small molecule addition alone. MED22 and INTS7 peptides were not identified in Experiment **A** therefore no Z-scores are presented. Strong stability shifts suggest small molecule driven changes in stability.

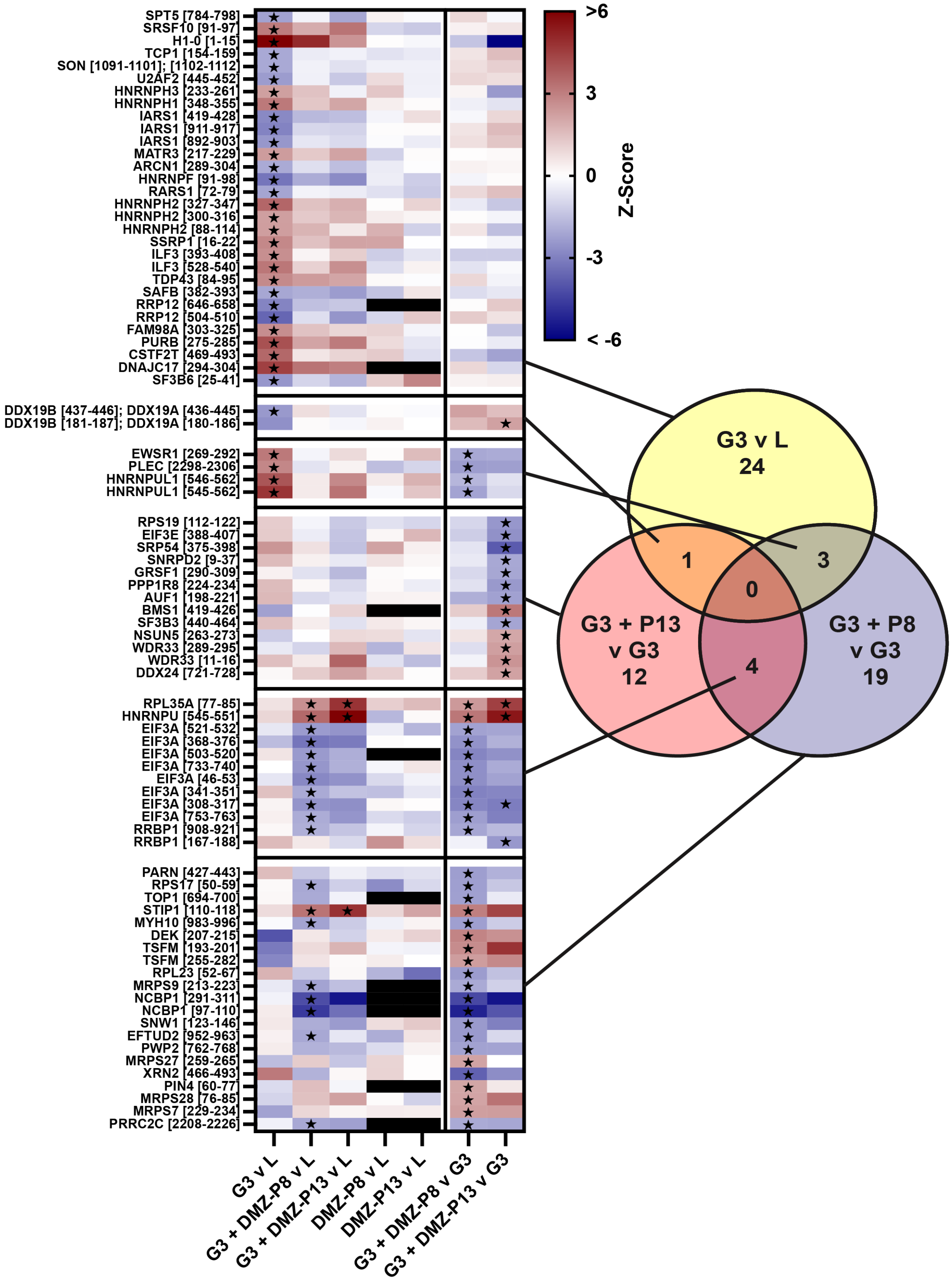


**Figure S10. Z-Score heatmap for all G3 v L and DMZ+G3 v G3 peptides.** All proteins and their corresponding peptides from **Figure 6** are shown with their Z-scores from all comparisons. Those highlighted with black stars correspond to the peptides called as hits (Z-score > +/- 2, p-value < 0.05) in those specific comparisons.

### Table S3. Hit DMZ structures, assay data and error.

| DMZ ID | Structures | IDA - $CD_{50} \pm$ SEM (μM) | | | | G3 DSF - $\Delta T_m \pm$ SD (°C) | | | G3 RT-qPCR - $\Delta C_t \pm$ SEM | |
|---|---|---|---|---|---|---|---|---|---|---|
| | | G1 | G2 | G3 | N1 | 5 μM | 25 μM | N | dCt | N |
| DMZ-P5 | 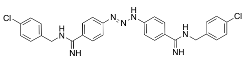 | 17.24 ± 12.3 | 5.12 ± 1.39 | 0.91 ± 0.13 | 2.07 ± 0.11 | 1.37 ± 0.34 | 3.24 ± 0.31 | 4 | -1.74 ± 0.18 | 5 |
| DMZ-P8 | 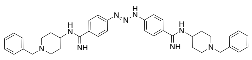 | 6.91 ± 0.46 | 5.12 ± 1.4 | 1.87 ± 0.08 | 15.5 ± 0.66 | 0.18 ± 0.24 | 2.00 ± 0.46 | 4 | 0.86 ± 0.17 | 7 |
| DMZ-P8Cl | 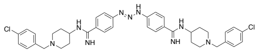 | 23.3 ± 2.8 | 33.5 ± 7.7 | 6.17 ± 0.27 | 10.3 ± 1.2 | 2.11 ± 0.42 | 4.40 ± 0.42 | 4 | -0.10 ± 0.12 | 3 |
| DMZ-P9 | 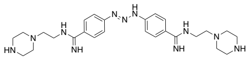 | 4.91 ± 2.21 | $25.8^{\perp}$ | 1.12 ± 0.49 | 9.69 ± 0.85 | 2.51 ± 0.26 | 5.28 ± 0.40 | 4 | -0.83 ± 0.20 | 4 |
| DMZ-P13 | 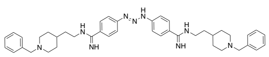 | 1.55 ± 0.20 | 1.91 ± 0.23 | 0.44 ± 0.26 | 4.02 ± 0.93 | 1.92 ± 0.28 | 3.66 ± 0.38 | 4 | -1.40 ± 0.18 | 5 |
| DMZ-P14 | 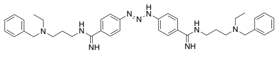 | 72.4 ± 20.7 | 30.5 ± 15.5 | 2.51 ± 0.37 | 13.2 ± 2.8 | 0.57 ± 0.32 | 2.62 ± 0.22 | 4 | -0.14 ± 0.24 | 3 |

| | | | | | | | | | | |
|---|---|---|---|---|---|---|---|---|---|---|
| DMZ-M1 | 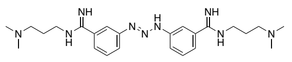 | 1.90 ± 0.04 | 3.55 ± 0.79 | 1.15 ± 0.18 | 5.29 ± 0.37 | *N/a | *N/a | 4 | -0.39 ± 0.23 | 4 |
| DMZ-M10 | 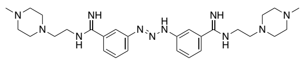 | 12.12 ± 2.36 | 14.46 ± 5.91 | 3.26 ± 0.31 | 48.6 ± 20.12 | 1.37 ± 0.34 | 3.24 ± 0.31 | 4 | -0.32 ± 0.29 | 4 |

⊥Only one replicate reported due to well inconsistency during screening.
*DMZ-M1 was not measured in DSF due to lack of melting peaks when DMZ-M1 was added.

#### Table S4. Stratifying SPROX-MS hits by RNA-binding proteins and GQBPs.

All assayed **G3 +/- DMZ** protein hits versus lysate control

| | G3 | G3 + DMZ-P8 | G3 + DMZ-P13 | DMZ-P8 | DMZ-P13 |
|---|---|---|---|---|---|
| **All** | 53 | 171 | 48 | 22 | 60 |
| **Stabilized** | 25 | 95 | 35 | 11 | 29 |
| **Destabilized** | 28 | 67 | 13 | 11 | 29 |
| **Hit Rate (%)** | 3.7% | 11.9% | 3.4% | 1.3% | 3.4% |

GO RNA-binding proteins for **G3 +/- DMZ** protein hits versus lysate control

| | G3 | G3 + DMZ-P8 | G3 + DMZ-P13 | DMZ-P8 | DMZ-P13 |
|---|---|---|---|---|---|
| **All** | 28 | 96 | 20 | 12 | 27 |
| **Stabilized** | 16 | 53 | 15 | 8 | 15 |
| **Destabilized** | 12 | 38 | 5 | 4 | 10 |
| **Hit Rate (%)** | 4.5% | 15.4% | 3.2% | 1.7% | 3.9% |

GQBP-filtered proteins for **G3 +/- DMZ** protein hits versus lysate control

| | G3 | G3 + DMZ-P8 | G3 + DMZ-P13 | DMZ-P8 | DMZ-P13 |
|---|---|---|---|---|---|
| **All** | 14 | 54 | 8 | 6 | 17 |
| **Stabilized** | 5 | 31 | 5 | 4 | 10 |
| **Destabilized** | 9 | 26 | 3 | 2 | 7 |

**G3 + DMZ-P8/DMZ-P13 v G3 protein hits**

Overlap with DMZ hits refers to overlap within these comparisons to **DMZ-P8 v L** and **DMZ-P13 v L** hits. The overall lack of overlap suggests that the majority of hits being identified for statistically significant changes in stability are therefore **G3 + DMZ** dependent. Hit rate (%) was calculated based on Experiment **A** protein numbers for each of these categories.

| **Proteins Assayed** | **All Assayed** | | **RNA-binding proteins** | | **GQBPs** | |
|---|---|---|---|---|---|---|
| **Treatment Comparisons** | **G3 + DMZ-P8 v G3** | **G3 + DMZ-P13 v G3** | **G3 + DMZ-P8 v G3** | **G3 + DMZ-P13 v G3** | **G3 + DMZ-P8 v G3** | **G3 + DMZ-P13 v G3** |
| **All** | **54** | **41** | **26** | **17** | **11** | **8** |
| **Stabilized** | **26** | **23** | **9** | **7** | **5** | **6** |
| **Destabilized** | **28** | **18** | **17** | **10** | **6** | **2** |
| **Hit Rate (%)** | **3.8%** | **2.9%** | **4.2%** | **2.7%** | **4.2%** | **3.0%** |

| | | | | | | |
|---|---|---|---|---|---|---|
| **Overlap with DMZ hits** | 1 | 4 | 0 | 1 | 0 | 1 |

**Table S5. Sequences and constructs used in this study.**

| ***Oligo Sequence Name*** | ***Nucleotide Sequence (5' to 3')*** |
|---|---|
| *DNA duplex template MALAT1 12bp-flanking* ***G1*** | Sense: GAA ATT AAT ACG ACT CAC TAT AGG CGG TGC TTG AAG GGG AGG GAA AGG GGG AAA GCG GGC AAC CAC TTT TC<br><br>Antisense: mGmAA AAG TGG TTG CCC GCT TTC CCC CTT TCC CTC CCC TTC AAG CAC CGC CTA TAG TGA GTC GTA TTA ATT TC |
| *DNA duplex template MALAT1 12bp-flanking* ***G2*** | Sense: GAA ATT AAT ACG ACT CAC TAT AGC TGG AAT TTG GAG GGA TGG GAG GAG GGG GTG GGG CTT ACT TGT TGT<br><br>Antisense: mAmCA ACA AGT AAG CCC CAC CCC CTC CTC CCA TCC CTC CAA ATT CCA GCT ATA GTG AGT CGT ATT AAT TTC |
| *DNA duplex template MALAT1 12bp-flanking* ***G3*** | Sense: GAA ATT AAT ACG ACT CAC TAT AGT GAC CTT ATA TAG GGA AGG GAG GGG GTG CCT GTG GGG TTT TAA AGA AT<br><br>Antisense: mAmUT CTT TAA AAC CCC ACA GGC ACC CCC TCC CTT CCC TAT ATA AGG TCA CTA TAG TGA GTC GTA TTA ATT TC |
| *DNA duplex template MALAT1 26bp-flanking* ***G3*** | Sense: GAA ATT AAT ACG ACT CAC TAT AGA TGT TTT ACA CTA TTG ACC TTA TAT AGG GAA GGG AGG GGG TGC CTG TGG GGT TTT AAA GAA TTT TCC TTT GCA GAG G<br><br>Antisense: mCmCT CTG CAA AGG AAA ATT CTT TAA AAC CCC ACA GGC ACC CCC TCC CTT CCC TAT ATA AGG TCA ATA GTG TAA AAC ATC TAT AGT GAG TCG TAT TAA TTT C |
| *RNA MALAT1 12bp-flanking* ***G1*** | GCG GUG CUU GAA GGG GAG GGA AAG GGG GAA AGC GGG CAA CCA CUU UUC |
| *RNA MALAT1 12bp-flanking* ***G2*** | CUG GAA UUU GGA GGG AUG GGA GGA GGG GGU GGG GCU UAC UUG UUG U |
| *RNA MALAT1 12bp-flanking* ***G3*** | UGA CCU UAU AUA GGG AAG GGA GGG GGU GCC UGU GGG GUU UUA AAG AAU |
| *RNA MALAT1 26bp-flanking* ***G3*** | ATG TTT TAC ACT ATT GAC CTT ATA TAG GGA AGG GAG GGG GTG CCT GTG GGG TTT TAA AGA ATT TTC CTT TGC AGA GG |
| *Forward primer 26bp-flanking* ***G3*** | ATA TAA GGT CAA TAG TGT AAA ACA T |

| | |
|---|---|
| *Reverse primer 26bp-flanking **G3*** | CCT CTG CAA AGG AAA ATT CTT TAA A |

**DMZ-P8Cl characterization and synthesis.**

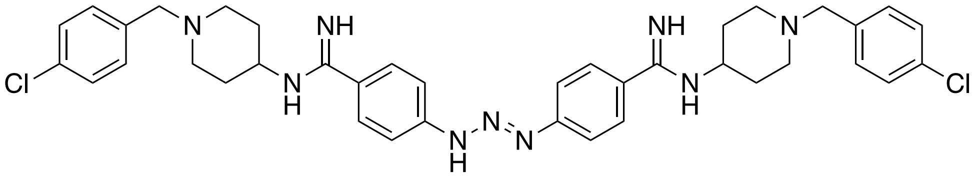


Bright orange powder, yield = 56%. $^{1}$H NMR (500 MHz, Methanol-d4) δ 7.78 – 7.75 (m, 6H), 7.68 – 7.65 (m, 3H), 7.63 – 7.60 (m, 3H), 7.36 (s, 6H), 3.58 (s, 4H), 3.01 – 2.97 (m, 3H), 2.24 (td, J = 11.9, 2.3 Hz, 3H), 2.09 – 2.05 (m, 3H), 1.82 – 1.75 (m, 3H). $^{13}$C NMR (126 MHz, MeOD) δ 163.18, 133.30, 130.66, 129.01, 128.03, 126.28, 118.56, 118.16, 117.90, 107.29, 61.49, 51.52, 50.60, 48.11, 30.19.

Synthesis of para-biscyano scaffold and substituent addition was performed as described in previous work.[2]
